# Transient Notch Activation Converts Pluripotent Stem Cell-Derived Cardiomyocytes Towards a Purkinje Fiber Fate

**DOI:** 10.1101/2024.09.22.614353

**Authors:** David M. Gonzalez, Rafael Dariolli, Julia Moyett, Stephanie Song, Bhavana Shewale, Jacqueline Bliley, Daniel Clarke, Avi Ma’ayan, Stacey Rentschler, Adam Feinberg, Eric Sobie, Nicole C. Dubois

## Abstract

Cardiac Purkinje fibers form the most distal part of the ventricular conduction system. They coordinate contraction and play a key role in ventricular arrhythmias. While many cardiac cell types can be generated from human pluripotent stem cells, methods to generate Purkinje fiber cells remain limited, hampering our understanding of Purkinje fiber biology and conduction system defects. To identify signaling pathways involved in Purkinje fiber formation, we analyzed single cell data from murine embryonic hearts and compared Purkinje fiber cells to trabecular cardiomyocytes. This identified several genes, processes, and signaling pathways putatively involved in cardiac conduction, including Notch signaling. We next tested whether Notch activation could convert human pluripotent stem cell-derived cardiomyocytes to Purkinje fiber cells. Following Notch activation, cardiomyocytes adopted an elongated morphology and displayed altered electrophysiological properties including increases in conduction velocity, spike slope, and action potential duration, all characteristic features of Purkinje fiber cells. RNA-sequencing demonstrated that Notch-activated cardiomyocytes undergo a sequential transcriptome shift, which included upregulation of key Purkinje fiber marker genes involved in fast conduction such as *SCN5A, HCN4 and ID2,* and downregulation of genes involved in contractile maturation. Correspondingly, we demonstrate that Notch-induced cardiomyocytes have decreased contractile force in bioengineered tissues compared to control cardiomyocytes. We next modified existing *in silico* models of human pluripotent stem cell-derived cardiomyocytes using our transcriptomic data and modeled the effect of several anti-arrhythmogenic drugs on action potential and calcium transient waveforms. Our models predicted that Purkinje fiber cells respond more strongly to dofetilide and amiodarone, while cardiomyocytes are more sensitive to treatment with nifedipine. We validated these findings *in vitro*, demonstrating that our new cell-specific *in vitro* model can be utilized to better understand human Purkinje fiber physiology and its relevance to disease.

## Main

The ventricular conduction system (VCS) plays a critical role in stimulating the contraction of the ventricular myocardium. These specialized cardiomyocytes (CMs) receive electrical impulses from the supraventricular portions of the cardiac conduction system (CCS), namely the atrioventricular (AV) and sinoatrial nodal (SAN) cells, and accelerate the speed of the electrical impulse across the large ventricular chambers. Due to the importance of this coordination in maintenance of hemodynamic stability, arrhythmias can result in serious consequences including sudden cardiac death (Haissaguerre et al., 2016). These arrhythmias may arise early in life as a result from improper positioning of the CCS during development, from mutations in genes responsible for normal development and function of the CCS (Baruteau et al., 2015; Bruneau and Srivastava, 2014), from fibrotic scarring following myocardial infarction, or from commonly prescribed medications which alter electrolyte imbalances critical for normal cardiac physiology.

Cellular and molecular studies to uncover the mechanisms underlying arrhythmias have been hampered by the inability to maintain various cardiomyocyte (CM) subtypes *ex vivo* in extended culture. Many of the cardiovascular cell types have been derived from human pluripotent stem cells (hPSCs) in recent years (Burridge et al., 2014; Calderon et al., 2016b; Chadwick et al., 2003; Iyer et al., 2015a; xref>; Kattman et al., 2011a; Kaufman et al., 2001; Lee et al., 2017b; Lian et al., 2012; Lian et al., 2013a; Mikryukov et al., 2021; Murry and Keller, 2008; Protze et al., 2017; Yang et al., 2016). These hPSC models have provided a greater understanding of cardiac physiology, serving as platforms for cell therapy and drug discovery. However, similar advances in the use and creation of human cardiac PFs have lagged behind. Until recently, protocols to generate cardiac PFs *in vitro* had only been reported using mouse PSCs (Maass et al., 2015; Tsai et al., 2015). More recently, *Prodan et al* identified a cocktail of several small molecules to reprogram hPSC-CMs toward a PF-like fate (Prodan et al., 2022). These studies offer a significant step toward use of PF-like cells in culture. However, a comprehensive molecular characterization of the cells resulting from these protocols is still lacking, as is their utility for a better understanding of differences between hPSC-derived CMs and PFs under physiological challenges that may be relevant to disease.

CMs and PFs share many characteristics and express similar markers but differ in key features concordant with their function. In comparison to CMs, PF have an elongated morphology and decreased sarcomere organization with decreased mitochondrial content and increased glycogen storage. In order to more rapidly propagate electrical impulses across the large ventricular chambers, PFs have faster conduction velocity and a longer action potential duration (APD) with a faster upstroke velocity (Park and Fishman, 2017; van Eif et al., 2018; van Weerd and Christoffels, 2016). PFs also express higher levels of fast-conduction markers such as CX40 and SCN5A. Recent studies profiling the mouse embryonic CCS at single-cell resolution have identified a number of novel markers of PF identity including *Cacna2d2, Igfbp5* and *Sema3a* (Goodyer et al., 2019). These data also indicated that expression of adult markers of PF identity such as *Cntn2* may not be fully recapitulated in embryonic PFs.

A possible explanation for the paucity of PF protocols may be that PF development is complex, and still poorly understood compared to that of other cells in the heart. PFs, unlike other cell types of the CCS, share a common progenitor with the ventricular working myocardium (Miquerol et al., 2010). Fate mapping studies in mice show a dual contribution of the trabecular myocardium to the PF network and the working myocardium, however the signaling pathways governing PF specification remain to be elucidated. The Notch signaling pathway is known to play several key roles during ventricular chamber development, including regulation of trabeculation, and CM proliferation and differentiation (D’Amato et al., 2016c; Luxán et al., 2016; MacGrogan et al., 2010; Zhao et al., 2019). Previous work showed that transient activation of Notch signaling can convert murine ventricular CMs toward a conductive-like fate (Khandekar et al., 2016; Rentschler et al., 2012), as characterized by changes in cardiac action potential (AP) morphology, increased expression of fast-conduction markers, and epigenetic changes to promoters and regulators of K+ channels.

In this study, we sought to develop a new method to generate and characterize human PSC-derived PF cells. Using publicly available single-cell RNA sequencing (scRNAseq) data from murine ventricles, we identified differences in Notch signaling between embryonic PFs and CMs. We show that transient Notch signaling activation in hPSC-CMs results in gradual changes toward a more elongated morphology and in changes of electrophysiological properties including an increased APD, spike slope and conduction velocity. These changes were accompanied with increased expression of fast-conduction markers. RNA sequencing of control and Notch activated hPSC-CMs uncovered differences in Notch and BMP signaling, as well as processes involved in the regulation of cardiac conduction, cell-cell communication and adhesion, and calcium handling, indicating acquisition of a PF-like fate. Utilizing transcriptional data from hPSC-derived CMs and PFs, we modified existing *in silico* excitation-contraction coupling models to simulate differences in response to various drugs and other physiological stimuli on action potential (AP) and calcium transient (CaT) waveforms. We identified several classes of drugs that demonstrated increased sensitivity in PFs compared to CMs and validated these findings experimentally. These studies demonstrate the utility of our new *in vitro* PF model to uncover aspects of human PF biology and their relevance to disease, and they illustrate the power of this model to serve as a platform for further drug discovery, which may have implications for the treatment and management of ventricular arrhythmias.

## Results

### Murine embryonic Purkinje fibers are enriched for Notch signaling components

To directly uncover the signaling pathways that discern early PFs from CMs, and thus might represent avenues to generate PFs *in vitro*, we examined published scRNA-seq data of E16.5 murine cardiac tissues that sought to identify new markers of the CCS lineages (Goodyer et al., 2019). Clustering and differential expression (DE) analysis of dissected subendocardial tissue identified several CM clusters as well as smooth muscle cells, fibroblasts and endothelial/endocardial cells **(Figure 1A-C)**. In order to focus on the CM populations we subclustered CMs, revealing additional heterogeneity within this population **(Figure 1D)**. We found that ventricular CMs separated broadly into compact and trabecular myocardium on the basis of *Hey2* and *Nppa* expression respectively **(Figure 1E)**. Similar to the published analysis we also identified a population that selectively expressed markers characteristic for PFs, such as *Cacna2d2, Scn5a, Id2* and *Gja5* among others **(Figure 1F)**.

**Figure 1.**
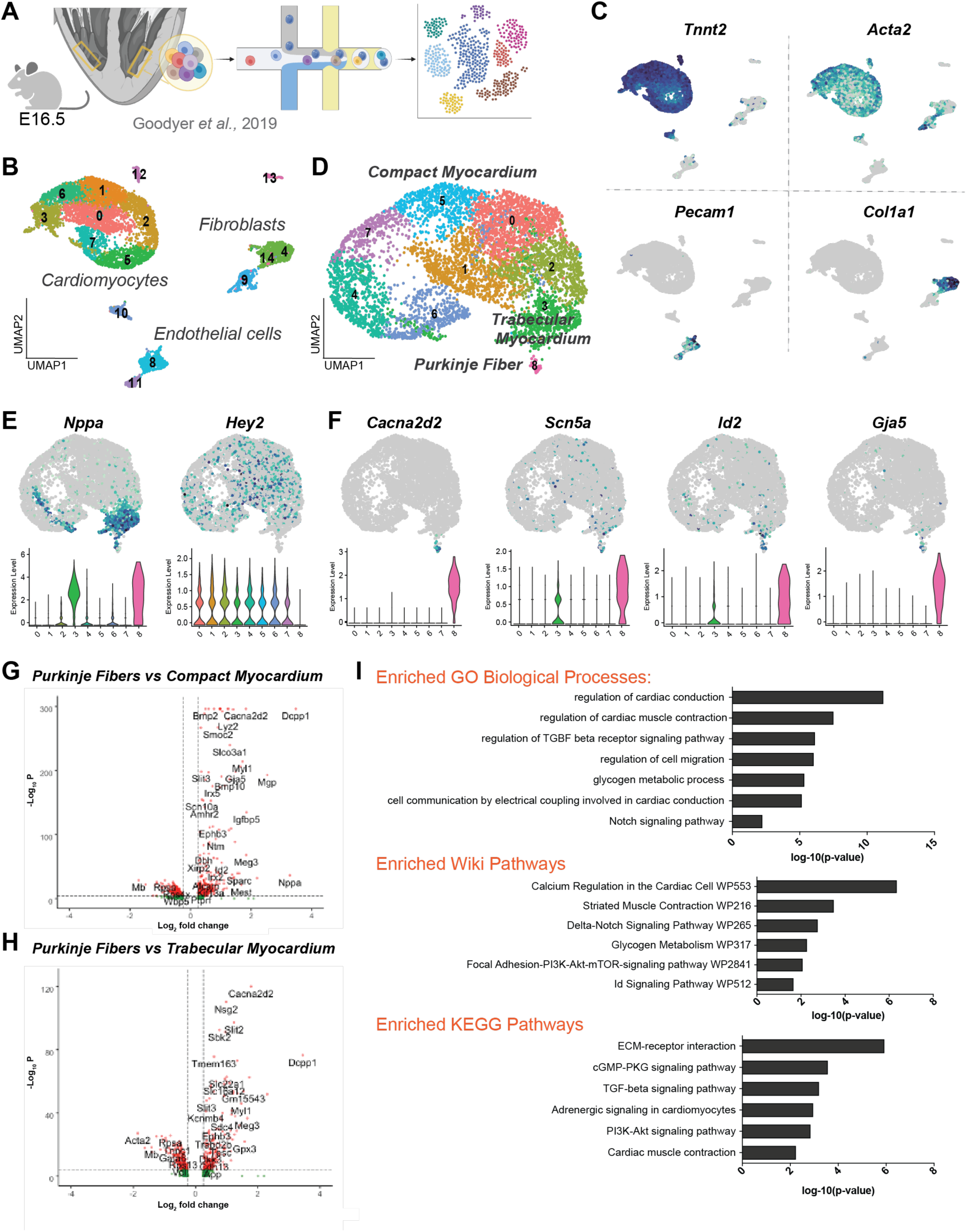
Analysis of murine embryonic cardiomyocyte and Purkinje fiber single-cell signature reveals differential expression of Notch signaling. **A)** *Top:* Schematic of experiment performed by (Goodyer et al., 2019) to obtain embryonic ventricular tissue dissected to enrich for bundle branch and Purkinje fiber cell types. **B)** UMAP projection of single cell sequencing data from Goodyer et al 2019 obtained from E16.5 embryonic mouse ventricular tissue. **C)** Feature plots for representative genes of major populations identified, including CMs (*Tnnt2+/Acta2+),* endothelial cell types (*Pecam1+*) and fibroblast populations (*Col1a1+*). **D)** UMAP projection following subclustering of CMs (top left of Figure 32B) identifies transcriptionally distinct PF cluster as well as CMs of trabecular and compact identity. **E/F)** Feature plots (top) and violin plots (bottom) for (E) compact/trabecular cardiomyocyte markers and (F) Purkinje fiber markers. **G)** Violin plot showing differentially expressed genes between PFs and compact myocardium. **H)** Violin plot showing differentially expressed genes between PFs and trabecular myocardium. **I)** GO (top), Wiki Pathway (middle) and KEGG Pathway (bottom) enrichment analysis for DE comparison between Purkinje fiber and trabecular myocardial cells demonstrating enriched terms and pathways.

We reasoned that by directly comparing PFs to ventricular CMs during the period of ventricular development we could identify genes and pathways that are involved in establishing PF identity. Toward this end we performed pairwise differential expression (DE) analysis between compact/trabecular CMs (clusters 1-7) and the PF population (cluster 8). As expected, this identified a number of aforementioned PF marker genes, but also members of signaling pathways such as *Bmp2, Bmp10, Ephb3,* and *Slit2/3* among others **(Figure 1G/H)**. We performed GO term and KEGG/Wiki Pathway enrichment analysis on these gene sets, identifying enriched processes such as glycogen metabolism, regulation of cardiac conduction and cell communication by electrical coupling, all consistent with a PF identity **(Figure 1I)**. Additionally, we uncovered enriched processes involved in Notch signaling in the PF cells.

### Transient activation of Notch signaling in hPSC-CMs results in changes in cellular morphology and marker expression characteristic of PF cells

Given that mouse PFs express Notch signaling components during development and that transient Notch activation has been shown reprogram mouse CMs into PF-like cells, we hypothesized that transient activation of Notch signaling could convert hPSC-CMs toward a PF-like fate. Toward this end, we used an hPSC line that ubiquitously expresses a Notch intracellular domain-Estrogen Receptor fusion protein (hereafter referred to as NICD-ER) (Ditadi et al., 2015). In this line, NICD-ER is ubiquitously expressed, and translocates to the nucleus upon the addition of 4-hydrodxytamoxifen (4OHT). To test our hypothesis, we treated hPSC-CMs with 4OHT for 1, 2, 3 and 7 days (d) to activate Notch signaling, or with the gamma secretase inhibitor DAPT to block endogenous Notch signaling, and then continued to culture the cells for four weeks **(Figure 2A)**. We found that hPSC-CMs treated with 4OHT for 2d gradually adopted a more elongated morphology, reminiscent of PFs, over the 4-week period. In contrast, hPSC-CMs treated with DAPT retained a cuboidal shape similar to untreated hPSC-CMs **(Figure 2B/C; Supplemental Figure 1)**. Treatment of hPSC-CMs with 4-OHT for 7d did caused cell death, suggesting that prolonged Notch activity is not compatible with hPSC-CM viability. As expected, the addition of 4OHT for 2d increased the expression of canonical Notch targets such as *HEY1, HEYL,* and *HES1,* whereas addition of DAPT reduced expression of these markers **(Figure 2D)**, confirming successful modulation of Notch signaling in hPSC-CMs.

**Figure 2.**
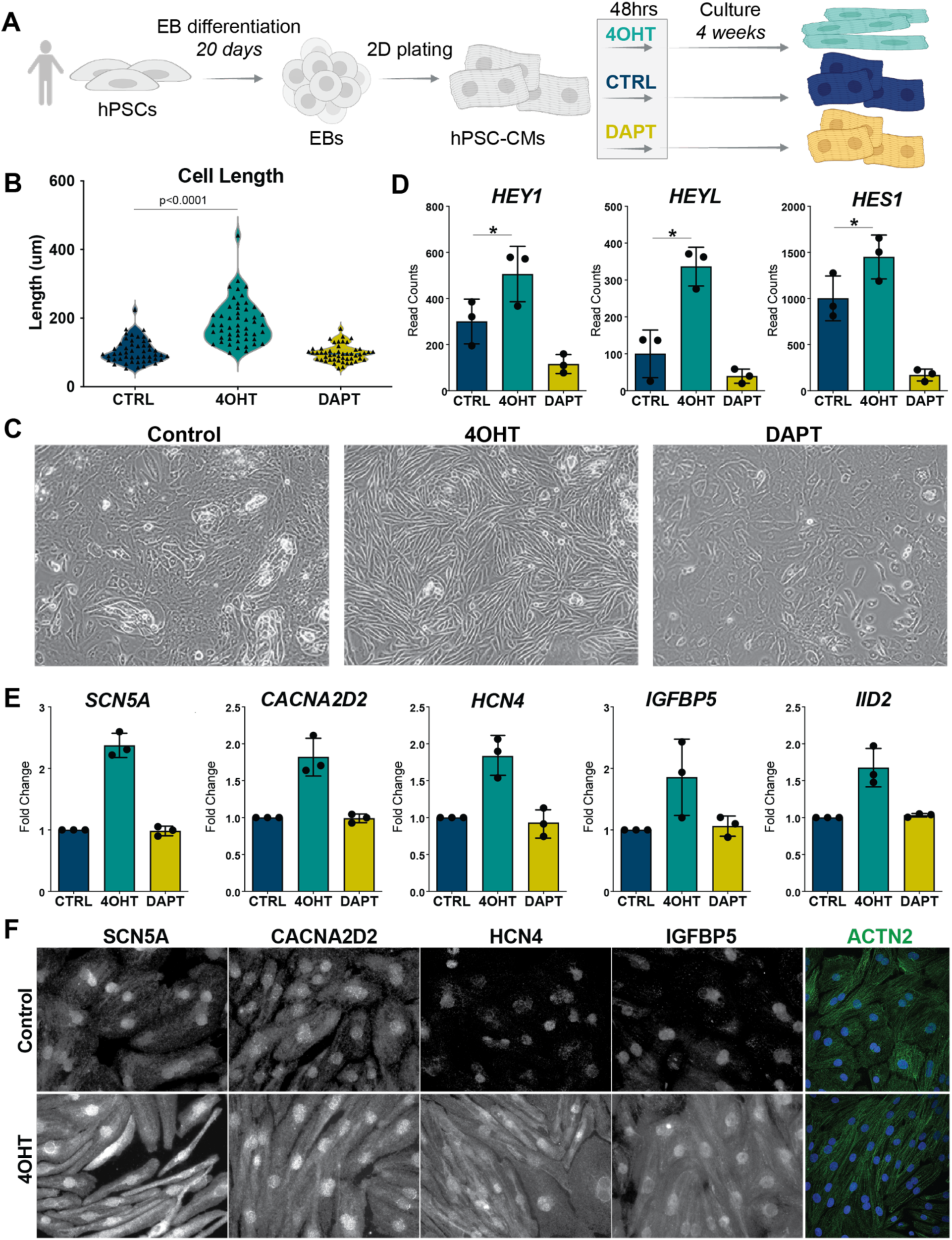
Transient Notch Upregulation in Pluripotent Stem Cell-Derived Cardiomyocytes Leads to Morphology Changes and Increased Expression of Purkinje Fiber Markers. **A)** Schematic demonstrating the strategy used to perturb Notch signaling in hPSC-CMs through addition of 4OHT or DAPT. **B)** Quantification of cell length between control and Notch perturbed hPSC-CMs. **C)** Treatment with 4OHT leads to development of elongated morphology of hPSC-CMs by day 28. **D)** RNA sequencing data demonstrating increased expression of canonical Notch targets 48 hours after induction/inhibition of Notch signaling in hPSC-CMs. **E)** RNA sequencing data demonstrating increased expression of PF markers in 4OHT induced hPSC-CMs 28 days after Notch perturbation. **F)** Immunofluorescence for alpha actinin (AACT - green) co-stained with DAPI (blue) and various PF marker genes (red) 28 days after Notch perturbation demonstrates increased expression of PF markers in 4OHT treated hPSC-CMs. Data is shown as fold change relative to time-matched replicate control. * = p <0.05 according to students t-test.

We performed RNA sequencing (RNA-seq) at day 28 (D28) and found that 4OHT-treated hPSC-CMs had increased mRNA expression of the PF markers *SCN5A, CACNA2D2, HCN4, IGFBP5,* and *ID2* **(Figure 2E)**. Immunofluorescence analysis confirmed that SCN5A, CACNA2D2, HCN4, and IGFBP5 were also elevated at the protein level **(Figure 2F)**. In contrast, DAPT-treated cells did not exhibit altered expression of these markers at either the mRNA or protein level. DAPT treatment also did not alter the sarcomere morphology of hPSC-CMs, as assessed by immunostaining for alpha actinin (ACTN2), indicating that endogenous Notch signaling is not required to maintain the CM identity of hPSC-CMs. These data suggest that transient Notch activation in hPSC-CMs caused stable changes in marker expression and cell shape, reminiscent of PF cells. For simplicity of data description we will subsequently refer to D28 4OHT-treated hPSC-CMs as hPSC-PFs.

### The global transcriptome of hPSC-PFs indicates a Purkinje fiber-like identity

In order to assess changes on the molecular level following Notch induction/inhibition, we performed principal component analysis (PCA) of the D28 RNA-seq data, which demonstrated that hPSC-PFs clustered away from untreated and DAPT-treated hPSC-CMs **(Figure 3A)**. This suggests that the global transcriptomes of hPSC-PFs and hPSC-CMs are distinct, potentially reflecting different cell identities. Specifically, we identified 465 DE genes between hPSC-PFs compared to untreated and DAPT-treated hPSC-CMs **(Figure 3B/C)**. As mentioned above, PF markers were upregulated in hPSC-PFs **(Figure 1E)**. The most downregulated genes included cell adhesion genes, such as *PCDH9*, which has previously been implicated in endothelial-mesenchymal transition during formation of the cardiac jelly and valve development, but does not have a known role in CMs (Feulner et al., 2022). A number of ADMTS family members were strongly downregulated, including *ADAMTS1*, *ADAMTS9* and *ADAMTS6*. Notably, ADAMTS1 regulates trabeculation initiation in the endocardium, but at later stages of development is involved in cessation of trabeculation due to closure of the cardiac jelly (Stankunas et al., 2008). Relatedly, a number of collagen family members such as *COL3A1*, *COL1A2*, *COL14A1*, and *COL12A1* were all downregulated in hPSC-PFs, suggesting that Notch activation alters the extracellular matrix.

**Figure 3.**
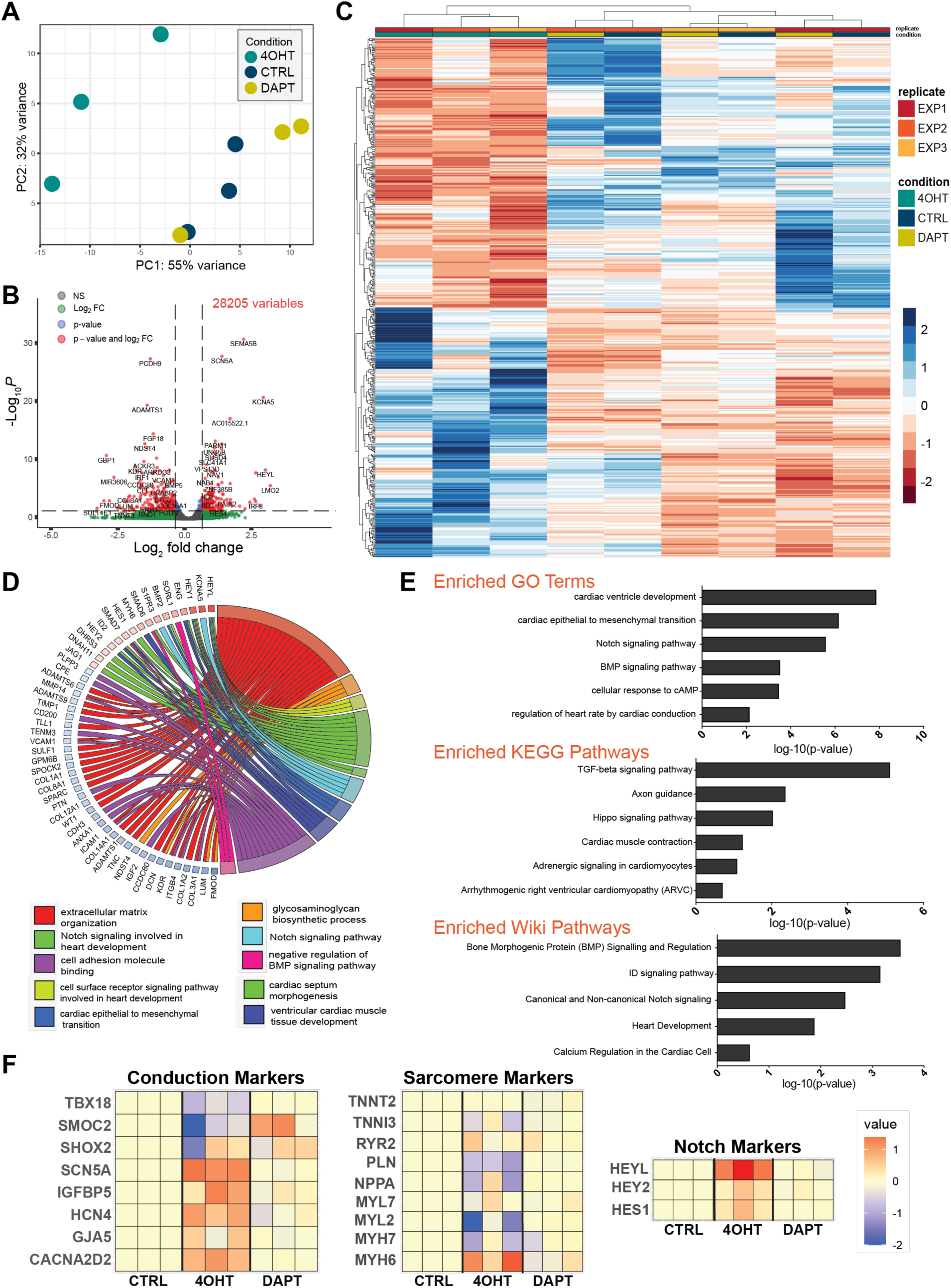
Transient Notch Activation in Pluripotent Stem Cell-Derived Cardiomyocytes Causes Global Transcriptomic Changes. **A)** Principal component analysis of bulk RNAseq data from control and Notch perturbed hPSC-CMs at d28. **B)** Volcano plot demonstrating results of **C)** Clustered heatmap demonstrating top 100 differentially expressed genes between control and 4OHT-treated hPSC-CMs. **D)** Chord plot demonstrating gene set enrichment analysis and GO processes that are associated with 4OHT-treated hPSC-CMs at d28. **D)** GO and KEGG/Wiki Pathway enrichment analysis on 4OHT-treated PSC-CMs at d28 demonstrating enrichment of processes involved in conduction and heart rhythm regulation. **F)** Heatmap demonstrating log2-fold change expression changes of selected genes at d28.

In order to uncover Notch-induced changes in biological processes and pathways, we performed GO and KEGG/Wiki Pathways enrichment analysis **(Figure 3D/E)**. We detected concomitant changes in a number of genes involved in BMP signaling and regulation, consistent with previous work implicating Notch signaling upstream of BMP and NRG/Ephrin signaling in the developing ventricle during trabeculation (D’Amato et al., 2016b; Grego-Bessa et al., 2007b). We also observed enrichment of genes involved in the epithelial-mesenchymal transition, consistent with the observed Notch-dependent changes in cell morphology. Importantly, we observed an enrichment for signals and processes that regulate heart rate and cardiac conduction, as well as arrhythmogenesis of the right ventricle, supporting a shift in cell type towards a more conductive fate. Moreover, pairwise DE analysis between embryonic PF and CM datasets (*Goodyer et al* 2019) and between hPSC-CMs and hPSC-PFs revealed 183 overlapping GO terms, including BMP signaling, regulation of cardiac contraction and adrenergic signaling **(Supplementary Figure 2A)**.

Next, we examined a number of candidates involved in cardiac conduction, sarcomere formation and Notch signaling. Notably, while fast conduction markers like *SCN5A, CACNA2D2* and *ID2* were upregulated, there was downregulation and/or no change in the expression of slow-conducting nodal markers such as *TBX18, SMOC2,* and *SHOX2* **(Figure 3F)**. From these data we infer that specialization towards conduction most resembled that of the fast-conducting ventricular conduction system. Compared to untreated and DAPT-treated hPSC-CMs, hPSC-PFs displayed a downregulation of genes involved in sarcomere organization and contractility, such as *PLN* and *MYL2,* and a lower *MYH7/MYH6* ratio, suggesting a less mature sarcomere phenotype. These data further support the conclusion that 4 weeks after transient Notch signaling induction, hPSC-PFs display a distinct transcriptional state resembling a PF-like cell identity.

### hPSC-CM reprogramming to hPSC-PF occurs sequentially and involves discrete changes in signaling pathways related to cardiac development

To further investigate the conversion of hPSC-CMs toward a PF-like fate, we performed RNA-seq at 1 day, 2 days, 3 days, 7 days, 14 days, and 28 days after 4OHT or DAPT treatment **(Figure 4A)**. We were intrigued to find that when comparing overlap of DE genes across these time points, the majority of the genes were differentially expressed at a single time point **(Figure 4B)**. A small subset of DE genes were shared between all the time points **(Figure 4B)**. This suggested to us that conversion toward a PF-like state involves a cascade of different genes that were being modulated sequentially, rather than a single transcriptional program that was being activated and maintained throughout the conversion process. In order to determine whether this extended to not only genes but particular processes, we performed GO enrichment at each stage and visualized the overlap of processes for the early (d1-3) and later time points (d7-28) **(Figure 4C)**. Similar to the DE genes, many processes were uniquely enriched at distinct stages, and several core processes were enriched across multiple time points. Amongst these, Notch signaling was enriched throughout the conversion, with early expression of the Notch target *JAG1* noticeable as early as d1 **(Figure 4D/E; Supplementary Figure 2B)**. Genes involved in “cell communication involved in cardiac conduction” and “bundle of His cell to Purkinje myocyte communication” were induced early, including *SCN5A* and *GJA5*, with some PF makers (*SCN5A*, *HCN4*, *NAV1*) showing a continuous increase across the 4 weeks *GJA5* **(Figure 4D/E)**.

**Figure 4.**
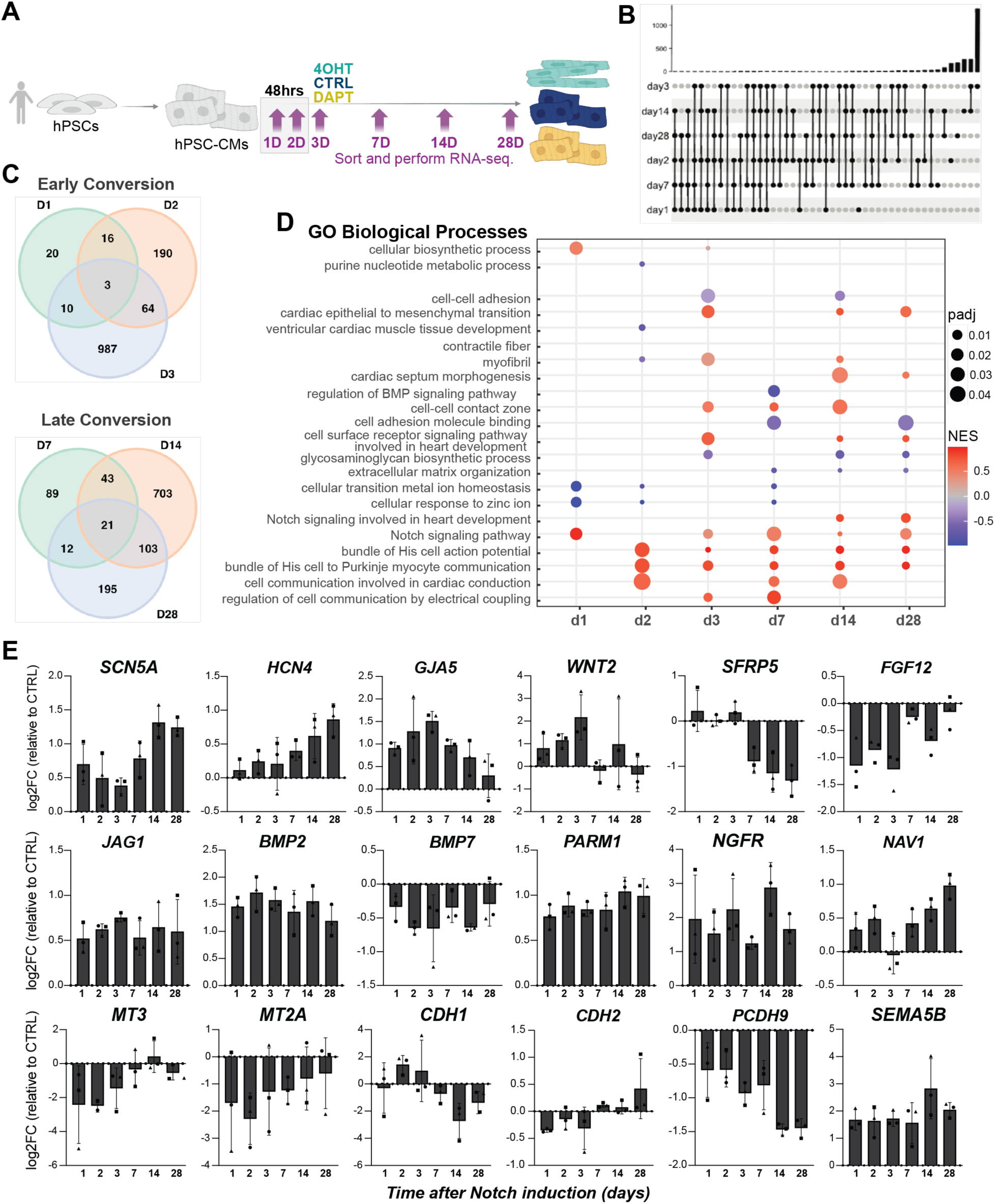
Conversion Toward Purkinje-like Lineage Following Notch Activation Occurs Through Gradual Expression of Stage Specific Genes and Processes. **A)** Schematic demonstrating strategy for converting hPSC-CM toward hPSC-PF lineage and time points collected for bulk RNAseq analysis. **B)** UpsetR plot demonstrating shared and exclusive differentially expressed genes between time points, demonstrating that most differentially expressed genes are exclusive to particular stages during conversion. **C)** Venn diagram demonstrating shared and exclusive GO terms between time points analyzed through bulk RNA sequencing. **D)** Bubble plot demonstrating expression of GO term enrichment at particular timepoints during conversion. **E)** Bar plots demonstrating log2-fold change expression of selected genes across conversion. Fold changes are calculated by normalizing gene expression in 4OHT treated cells to their experimental and time-matched control.

The early time points during Notch induction (d1 and d2) were also characterized by downregulation of processes involved in zinc ion homeostasis and cation trafficking, with rapid downregulation of a number of zinc transporters and metallothionein genes such as *MT3* and *MT2A* **(Figure 4D/E)**. Intriguingly, zinc transporters are essential for cardiac ventricular compaction and regulate the expression of metallothionein factors (Lin and Li, 2018; Lin et al., 2018). Downregulation of these genes is involved in increased glycosaminoglycan synthesis in the trabecular myocardium, and downregulation of ADAMTS family member enzymes (Lin and Li, 2018). These data are consistent with the suppression of a mature CM phenotype associated with ventricular compaction.

We also observed early changes in components of multiple signaling pathways that interact with Notch. For instance, *BMP2* and *PARM1* were upregulated and are involved in BMP/Smad signaling during CM development (Nakanishi et al., 2012)**(Figure 4E)**. Wnt family members such as *WNT2* and *SFRP5* were differentially expressed in hPSC-PFs, both of which have been shown to play key roles during ventricular development. Notch and Wnt signaling regulate CM proliferation, with Notch augmenting the expression of secreted Wnt agonists that boost CM proliferation (Zhao et al., 2019). Lastly, we observed a number of genes involved in epithelial to mesenchymal transition, with changes in cell-cell adhesion such as downregulation of *CDH1* and increased expression of processes involved in neural cell and neural crest migration (*GRID1, NGFR,* and *NAV1).* Among these was *SEMA5B,* a related semaphorin family member recently identified in mice as a specific marker for PF cells during development (Li et al., 2018). For any additional exploration of these data, we created an interactive and open access website (https://maayanlab.cloud/dubois-2/).

Overall, the temporal gene expression studies suggest that hPSC-CM conversion toward a PF-like fate involves sequential changes in gene expression and biological processes that resemble aspects of ventricular development and trabeculation initiation and maintenance. We propose that Notch activation in hPSC-CMs results in relative de-differentiation of hPSC-CMs and a subsequent promotion of a PF-like identity.

### hPSC-PFs have electrophysiological and contractile properties resembling *bona fide* PFs

CMs and PFs display distinct electrophysiological properties. Specifically, PFs have a higher upstroke velocity and increased conduction velocity compared to CMs. We observed a higher beat frequency for hPSC-PFs compared to untreated and DAPT-treated hPSC-CMs **(Figure 5A)**, consistent with increased expression of the cardiac conduction pathway and fast conduction markers such as HCN4 **(Figure 3E/F)**. To further investigate the electrophysiological properties, we cultured hPSC-CMs and hPSC-PFs on multi-electrode arrays (MEA), and measured their electrophysiological profile. We observed a decreased beat period and increased conduction velocity in hPSC-PFs compared to untreated and DAPT-treated hPSC-CMs **(Figure 5B/C)**. The spike slope, which reflects how fast Na+ ions move away from the electrode and inside the cell during the cardiac action potential (AP), was slower in hPSC-PFs compared to untreated and DAPT-treated hPSC-CMs, whereas the spike amplitude was higher.

**Figure 5.**
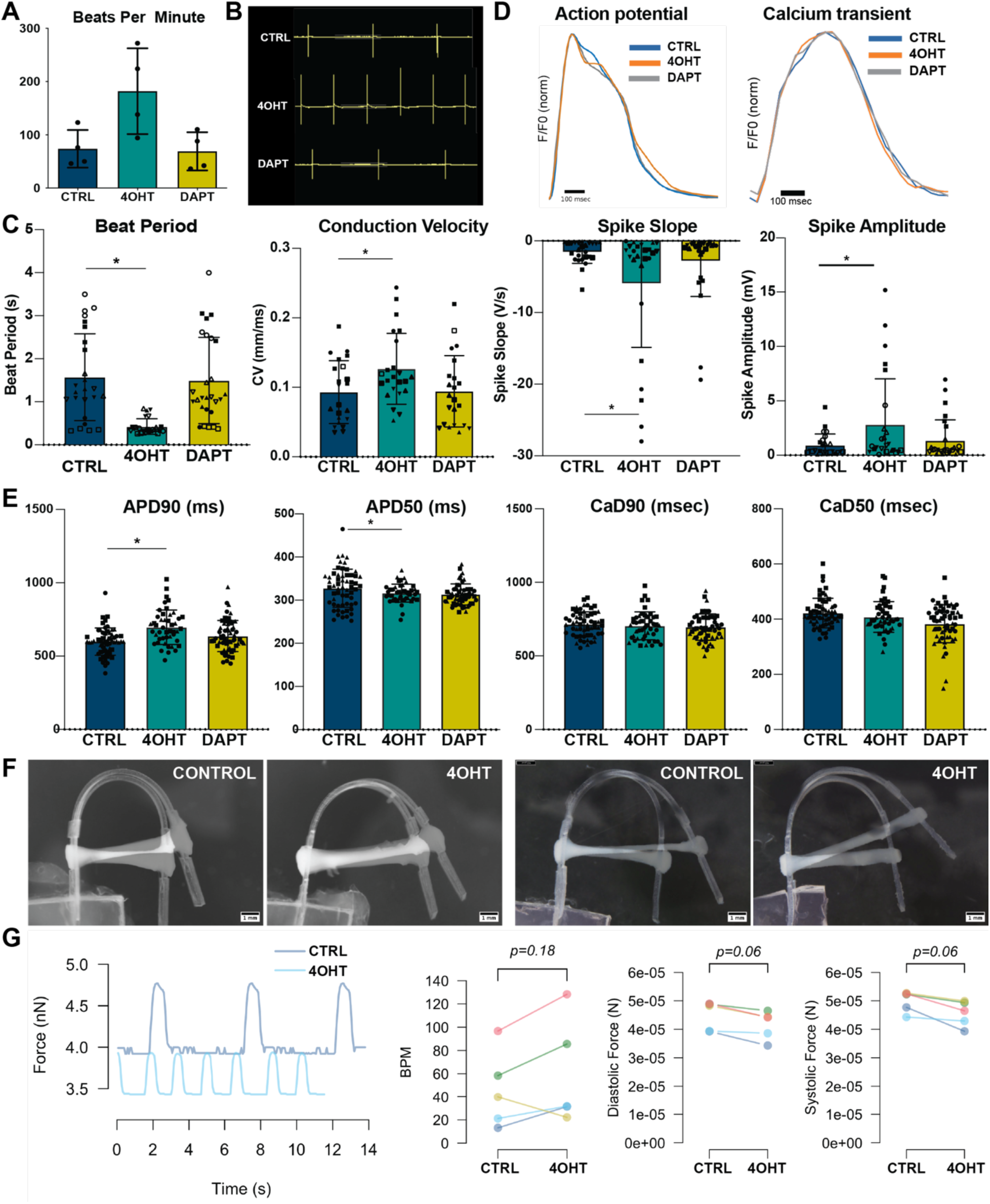
Transient Notch Activation in Pluripotent Stem Cell-Derived Cardiomyocytes Leads to Action Potential Morphology Changes. **A)** Quantification of changes in beat rate (manually counted) 28 days after removal of 4OHT or DAPT, as indicated. **B)** Representative traces of MEA recordings for hPSC-CMs in the indicated conditions. **C)** Quantification of beat period, conduction velocity, spike slope and spike amplitude in d28 hPSC-CMs corresponding to the indicated conditions. **D)** Representative traces of AP morphology (left) and of CaT morphology (right) analyzed using fluorescent dye for control and Notch perturbed hPSC-CMs. **E)** Quantification of APD90 and APD50 at d28, demonstrating increase in APD90 in Notch activated hPSC-CMs (left) and quantification of CaD90 and APD50 (right) at d28. **F)** Still images of N=2 representative dynEHT from hPSC-CMs and hPSC-PFs 4 weeks after casting. **G)** Force traces of hPSC-CM and hPSC-PF dynEHTs 4 weeks after casting (left) and analysis of beat per minute (BPM), diastolic force and systolic force (right) for N=5 replicates/differentiations. * = p <0.05 according to students t-test. Data consists of n=5 biological replicates, with 2-3 technical replicates per condition across each biological replicate.

Next, we simultaneously profiled the AP and calcium transient (CaT) features on a single-cell level, using a dual fluorescent dye-based method **(Figure 5D)**. The AP duration at 90% repolarization (APD90) was increased in hPSC-PFs compared to untreated and DAPT-treated hPSC-CMs, a hallmark difference between CM and PFs in mouse models, whereas the APD50 and CaT duration were unchanged **(Figure 5E)**.

Although the AP profiles of CMs and PFs have been compared in animal models (Árpádffy-Lovas et al., 2021; Han et al., 2000; Kristóf et al., 2012), there is a paucity of direct comparisons of human CMs and PFs. In order to better contextualize our findings in the context of human CMs and PFs, we took advantage of published computational models of these cell types that were established by transcriptomic and electrophysiological profiling of healthy adult human ventricular CMs and PFs (Tomek et al., 2019; Trovato et al., 2020). We simulated several AP and CaT waveforms in each model, obtained average waveform profiles using custom Matlab scripts, and overlaid them to directly compare the two cell types **(Supplemental Figure 3A/B, E/F)**. We found that, consistent with our findings using hPSC-CMs/hPSC-PFs, the APD90 was increased in PFs compared to CMs **(Supplemental Figure 3C/G/D**), though the magnitude of the increase in total APD was less than that previously reported in animal models (Iyer et al., 2015b; Khandekar et al., 2016). The models also revealed that, although CaTs had a slower upstroke in PFs compared to CMs **(Supplemental Figure 3H)**, there was no difference in the total duration of the CaT. Together, these observations suggest that the electrophysiological properties of hPSC-PFs are reminiscent of human PF cells, and that computational modeling confirms cell type-specific AP features of hPSC-CMs and hPSC-PFs found in adult CMs and PFs.

The main purpose of PFs is to rapidly propagate conduction across the ventricular tissue. PFs are contractile, albeit less than working cardiomyocytes. To test if this difference in contractility is recapitulated in hPAC-CMs/hPSC-PFs, we employed a dynamic tissue engineering (dynEHT) system. Both hPSC-CMs and hPSC-PFs were casted into dynEHTs, allowed to compact and measured 4 weeks after casting (**Figure 5F, Supplementary Videos 1-4**). Our analysis again confirmed the faster beat rate of hPSC-PFs, and also showed a decrease in both diastolic and systolic force in hPSC-PF dynEHTs compared to hPSC-CMs dynEHTs (**Figure 5G**).

### Computational models accurately predict distinct responses of hPSC-CM and hPSC-PF to drug exposure and electrolyte perturbations

Having established a cell type-specific hPSC model of CMs and PFs, we next explored the utility of this system to predict and uncover differential responses to biophysical stimuli, such as perturbations in ion conductance through channels commonly targeted by pharmacologic agents. To perform modeling predictions in hPSC-CMs and hPSC-PFs, we modified an *in silico* model of hPSC-CMs (Paci et al., 2018). From our bulk RNA-seq data on hPSC-CMs and hPSC-PFs we extracted genes corresponding to ion channels, pumps and transporters included in the Paci model. We calculated the fold changes in the expression of these genes and used these data to scale parameters in the Paci model to produce the hPSC-PF variant, taking into account differences in library size and depth of sequencing. The simulated AP profiles for hPSC-CMs and hPSC-PFs using this scaled computational model showed an increased duration of the AP in hPSC-PFs **(Figure 6A)**, similar to what we observed *in vitro* **(Figure 5E)**. Moreover, hPSC-PFs were predicted to have an increased beat frequency under spontaneous beating conditions **(Figure 6B)**, once again recapitulating our findings *in vitro* **(Figure 5A).** These data suggest that the scaled model accurately predicts key biological features of hPSC-CMs and hPSC-PFs.

**Figure 6.**
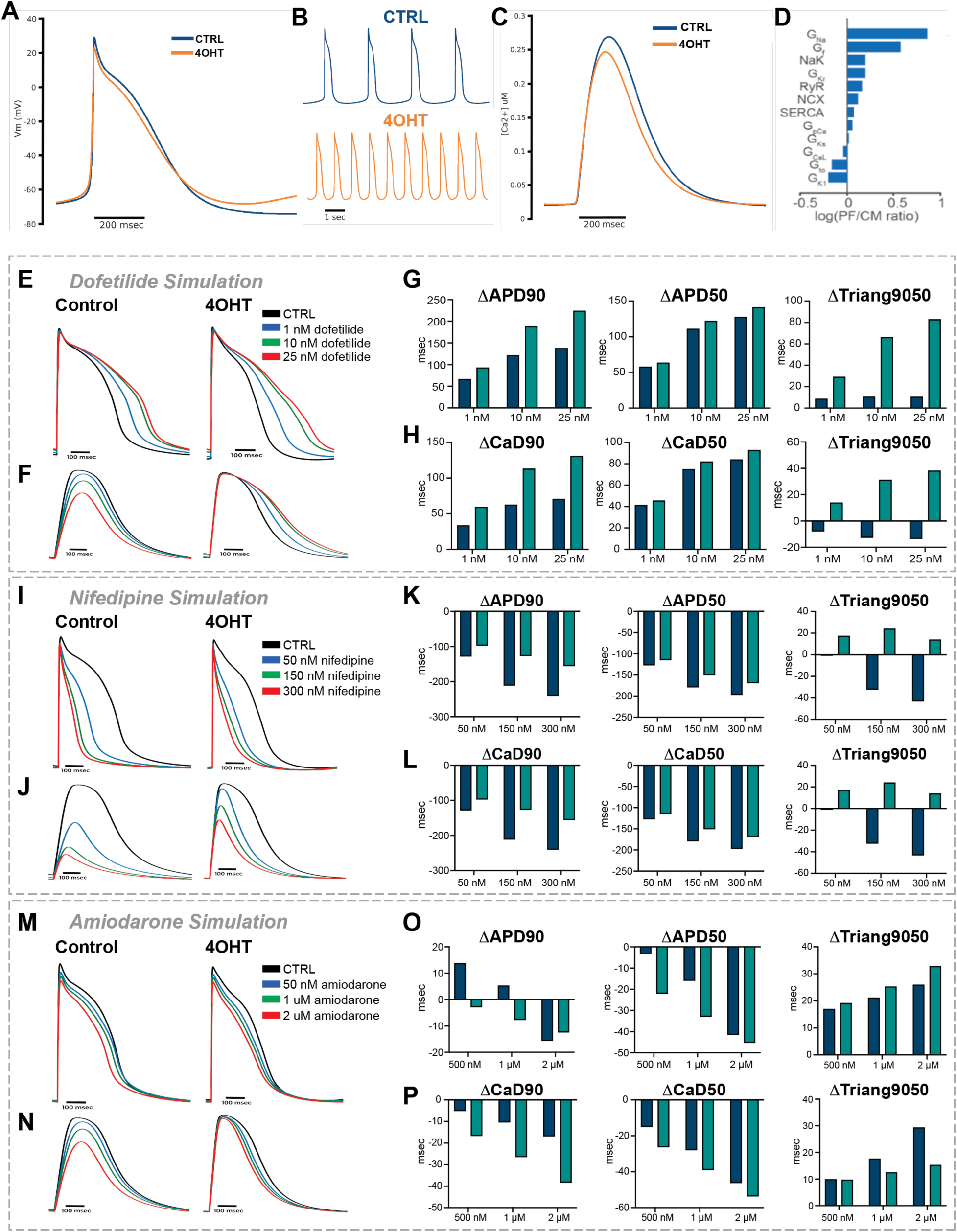
Computational Modeling of Pluripotent Stem Cell-Derived Cardiomyocytes and Purkinje Fibers Demonstrates Cell-Specific Sensitivity to Drug Perturbations. **A)** Simulated AP profile for hPSC-CMs (blue) and hPSC-PFs (orange) using a scaled computational model. **B)** Predicted spontaneous beating rate for hPSC-CMs (blue) and hPSC-PFs (orange). **C)** Predicted calcium transient profile for hPSC-CMs (blue) and hPSC-PFs (orange). **D)** Partial least squares regression analysis comparing the sensitivity of response to inhibition of commonly expressed channel proteins involved in CM physiology. **E)** Model simulation of the effect of dofetilide on the hPSC-CM/PF AP profile. **F)** Model simulation of the effect of dofetilide on the hPSC-CM/PF CaT profile. **G)** Quantification of the change in AP features across hPSC-CMs (blue) and hPSC-PFs (orange) as a function of dofetilide treatment. **H)** Quantification of the change in CaT features across hPSC-CMs (blue) and hPSC-PFs (orange) as a function of nifedipine treatment. **I)** Model simulation of the effect of nifedipine on the hPSC-CM/PF AP profile. **J)** Model simulation of the effect of nifedipine on hPSC-CM/PF CaT profile. **K)** Quantification of change in AP features across hPSC-CMs (blue) and hPSC-PFs (orange) as a function of nifedipine treatment. **L)** Quantification of the change in CaT features across hPSC-CMs (blue) and hPSC-PFs (orange) as a function of nifedipine treatment. **M)** Model simulation of the effect of modifying extracellular sodium concentration on hPSC-CM/PF AP profile. **N)** Model simulation of modifying extracellular sodium concentration on hPSC-CM/PF CaT profile. **O)** Quantification of the change in AP features across hPSC-CMs (blue) and hPSC-PFs (orange) as a function of sodium concentration changes. **P)** Quantification of the change in CaT features across hPSC-CMs (blue) and hPSC-PFs (orange) as a function of sodium concentration perturbation.

To identify differential sensitivity to perturbations in various ion channels across hPSC-CMs and hPSC-PFs, we performed a partial least squares regression analysis. Our modeling predicted that the APD in hPSC-PFs is more sensitive to perturbations of L-type Na channel, the chronotropic funny current (I_f_) and K+ inward rectifier currents (I_Kr_), whereas hPSC-CMs were predicted to be more sensitive to perturbations in L-type Ca2+ conductance **(Figure 6C/D)**.

We next extended these analyses to model the effect of commonly used drug classes that were predicted to target channels with cell-type sensitivity. The inward rectifier K+ channel inhibitor dofetilide was predicted to increase the APD and the CaT duration in both hPSC-CMs and hPSC-PFs as expected **(Figure 6E/F)**. However, the magnitude of the change in the APD50 and APD90 was predicted to be greater for hPSC-PFs than for hPSC-CMs **(Figure 6G)**. The magnitude of the change in the CaT duration at 90% repolarization was also predicted to be greater for hPSC-PFs than for hPSC-CMs **(Figure 6H)**. Dofetilide also increased the triangulation value (Triang9050) in hPSC-PFs compared to hPSC-CMs, a proxy which can be used as a predictor for risk of development of EAD/DADs (**Figure 6G/H**).

To determine if these simulations accurately predicted the responses to dofetilide experimentally, we treated hPSC-CMs and hPSC-PFs with and without dofetilide and recorded the AP and CaT waveforms using the dual fluorescence based optical system. As predicted by our models, we observed a prolonged APD and CaT in both hPSC-CMs and hPSC-PFs following treatment **(Figure 7A/B)**. Moreover, the magnitudes of the changes in the APD50, APD90, CaD50 and CaD90 were greater for hPSC-PFs than for hPSC-CMs **(Figure 7C-F)**, again consistent with our predictions. Notably, the predicted and observed magnitudes of the changes were slightly different, which may reflect distinct behaviors of hPSC-based cell types compared to the base model used in our computational simulations, or limitations of the inherent noise in the optical-based approach.

**Figure 7.**
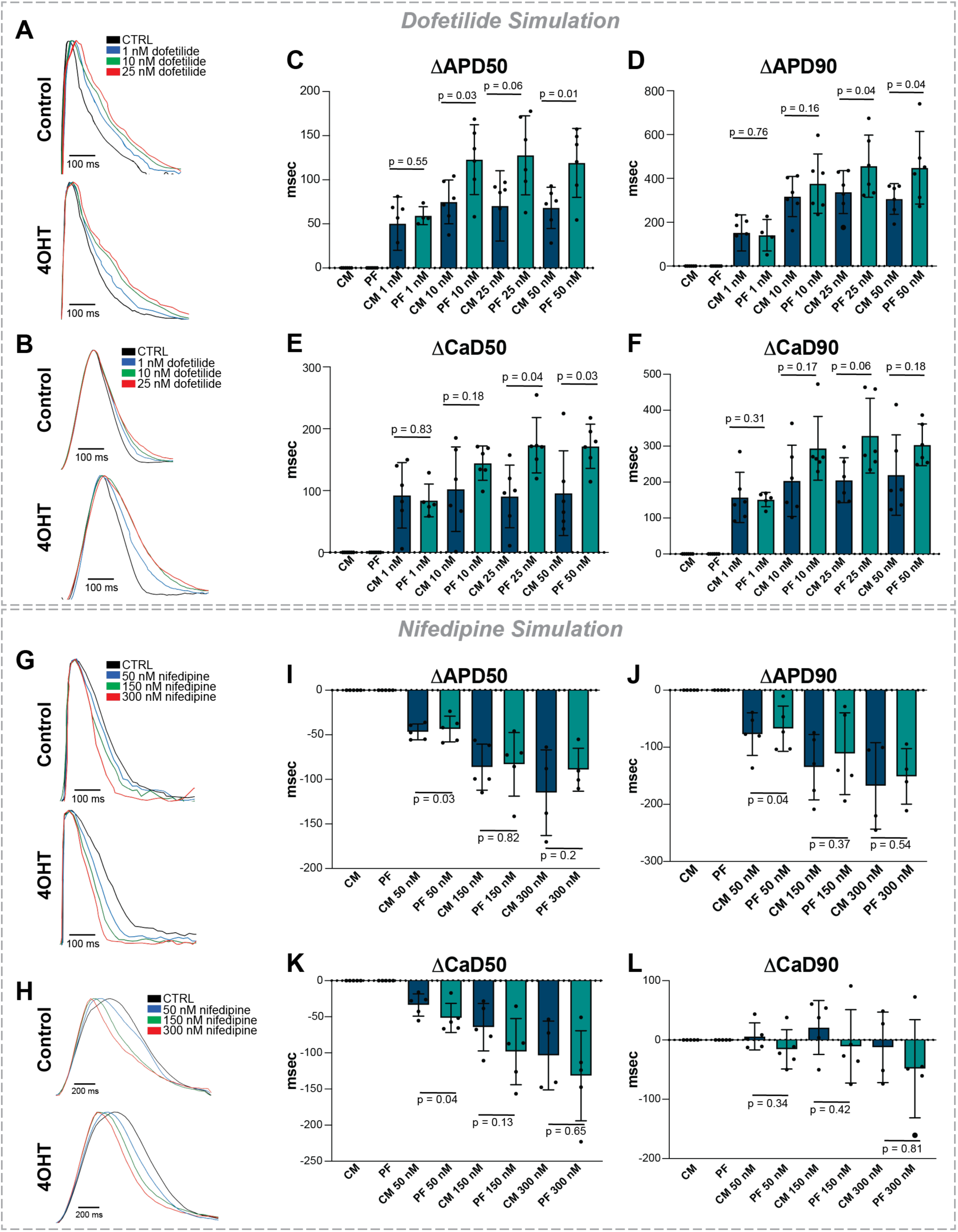
Pluripotent Stem Cell-Derived Purkinje Fibers Demonstrate Differential Sensitivity to Dofetilide and Nifedipine Treatment A/B) Representative (A) action potential and (B) CaT waveforms for hPSC-CMs and hPSC-PFs treated with varying concentrations of dofetilide. **C/D)** Quantification of changes in (C) APD50 and (D) APD90 in hPSC-CMs (blue) and hPSC-PFs (teal) compared to untreated controls. **E/F)** Quantification of changes in (E) CaD50 and (F) CaD90 in hPSC-CMs (blue) and hPSC-PFs (teal) compared to untreated controls. **G/J)** Representative (G) action potential and (J) CaT waveforms for hPSC-CMs and hPSC-PFs treated with varying concentrations of nifedipine. **H/I)** Quantification of changes due to nifedipine treatment in (H) APD50 and (I) APD90 in hPSC-CMs (blue) and hPSC-PFs (teal) compared to untreated controls. **K/L)** Quantification of changes in (K) CaD50 and (L) CaD90 in hPSC-CMs (blue) and hPSC-PFs (teal) compared to untreated controls. * = p <0.05 according to pairwise students t-test. Dots represent average of (n=10-15) technical replicates for each biological replicates for a total of (n=5) biological replicates.

We next looked at nifedipine, an L-type calcium channel inhibitor used to treat high blood pressure and arrhythmia. We modeled the effect of nifedipine on the AP and CaT waveforms and found that it decreased the APD50 and APD90 of both cell types in a concentration-dependent manner (**Figure 6I/K**). The magnitude of the change in APD was higher for hPSC-CMs compared to hPSC-PFs **(Figure 6I/K)**, consistent with previously published animal models (Terrar et al., 2007). Interestingly, while both cell types responded to nifedipine by decreasing the CaD50 and CaD90, hPSC-PFs were predicted to have a greater change in their CaD90 than hPSC-CMs **(Figure 6J/L)**. Thus, our computational models predict cell type-specific and AP/CaT-specific responses to nifedipine in human cells. To examine the responses to nifedipine *in vitro*, we treated hPSC-CMs and hPSC-PFs at d28 with increasing concentrations of nifedipine. Consistent with predictions from our model, we observed shortening of the APD and CaT in both hPSC-CMs and hPSC-PFs **(Figure 7G/J)** and hPSC-CMs displayed increased sensitivity to perturbation with nifedipine, demonstrating a greater magnitude of shortening in the APD50 and APD90 compared to hPSC-PFs **(Figure 7H/I)**. Moreover, compared to hPSC-CMs, hPSC-PFs exhibited an increased shortening of the CaD50 in response to nifedipine **(Figure 7K/L)**, similar to the trends observed in the model predictions, particularly of CaD90.

To extend our analyses to other physiological challenges, we investigated how hPSC-CMs and hPSC-PFs would respond to changes in extracellular ion concentrations. Specifically, we modeled the effect of low sodium (hyponatremic) and high sodium (hypernatremic) conditions. In both conditions, the predicted changes in APD and CaD were higher in magnitude for hPSC-CMs than hPSC-PFs **(Figure 6M/O)**. Through use of these models, we were able to simulate the effect of a large variety of drugs and other modulators of cardiac physiology, such as changing in concentrations of key ions to find “high-sensitivity” candidates that cause strong responses in a cell type-specific manner. These data are provided as a resource for further exploration **(Supplemental File S1).**

In summary, we find that hPSC-CMs and the newly established hPSC-PFs show distinct responses to common anti-arrhythmic medications, allowing for such analyses to be performed in a cell type-specific manner. We further show that computational models can accurately predict these distinct responses in a highly scalable manner. Overall, we demonstrate the utility of these cell type-specific models for studying regulators of cardiac physiology and for drug discovery.

## DISCUSSION

A thorough understanding of the molecular and cellular mechanisms underlying PF biology and the role of PF cells in ventricular arrhythmias is still needed. Progress toward that goal has been hampered previously by a lack of tools to study human PFs, which has implications not only in understanding disease pathology but also in advancing our understanding of cardiomyocyte-Purkinje cell interactions during homeostasis. In an effort to bridge that gap, we established that transient activation of Notch signaling can reprogram hPSC-CMs toward a PF-like fate characterized by changes in morphology, expression of key PF markers, and most importantly functional changes in electrophysiological properties that confer increased conductive ability. This novel cell-specific system will enable future interrogation of fundamental questions about PF biology, as well as empower pharmacological studies about cell-type specific differences in CM/PF responses to common physiological and/or genetic challenges.

Our DE analysis across multiple time points following Notch activation revealed discrete changes in specific genes at each time point with little overlap of core genes and biological processes across all time points. This suggests that a series of sequential events occurred downstream of Notch activation to direct hPSC-CMs toward an hPSC-PF identity. One of the few genes and processes that was involved at all time points was Notch signaling, which was unexpected given the transient nature of Notch activation in our protocol. We observed constitutive activation of Notch target genes such as *HEY1* and *HES1* several weeks after 4OHT treatment. This may suggest an underlying epigenetic mechanism which could reinforce Notch signaling in a more permanent manner. This would be consistent with previous studies in the mouse heart showing changes in promoter methylation status following Notch activation (Khandekar et al., 2016).

Relatedly, we found that while a number of PF markers such as *SCN5A* and *HCN4* were differentially regulated as early as d1 after Notch activation, these genes were more highly differentially expressed at d14 and d28. This may suggest that full acquisition of a PF-like identity is not an immediate consequence of Notch activation, and may involve additional downstream signaling pathways and mediators in a sequential fashion. We did observe that activation of BMP2 signaling occurred early after Notch activation, and remained differentially upregulated throughout. This was not entirely surprising given the well-established interactions between Notch and Ephrin/BMP signaling in the developing ventricle during trabeculation, and raises question of whether activation of BMP signaling may be necessary and sufficient downstream of Notch in the conversion process (Grego-Bessa et al., 2007b). It also raises questions about whether conversion of hPSC-CMs toward a PF-like identity through Notch activation represents activation of an endogenous developmental pathway or a synthetic programming strategy. Our analysis of published scRNA-seq studies of the developing mouse ventricle indicated that Notch signaling was enriched in embryonic PFs compared to trabecular progenitors. However, this was not the only signaling pathway that was enriched, and this analysis alone is not sufficient to conclusively to determine whether Notch plays a role in specifying the PF lineage. Further work profiling changes in the signaling environment of PFs across multiple stages of murine ventricular development would be required to more conclusively determine a potential role for Notch in endogenous generation of PF cell types during normal development.

Lastly, we demonstrated that the use of mathematical models in combination with our cell-type specific hPSC models offers tremendous utility for testing of cell-type specific responses in different cell type. Our models and subsequent *in vitro* validation experiments demonstrated for the first time that cell type-specific responses to drugs observed in other animal models could be recapitulated in human cell types (Árpádffy-Lovas et al., 2021; Terrar et al., 2007). This not only further underscores the biological similarity of hPSC-PFs to native PFs, but also serves as a proof-of-concept that these models can be used for further drug discovery of responses to other drug classes or other biophysical stimuli. The models established here allowed us to test representative examples of multiple drugs that have not previously been profiled in human CMs and PFs (or in select cases, in any model system). Additionally, the modeling studies included herein focused on the use of hPSC-based *in silico* models that best represent our cell types in culture. These analyses could be extended to take advantage of previously published adult CM/PF post-mortem derived models used earlier to benchmark differences in APD and CaD between CMs and PFs (Tomek et al., 2019; Trovato et al., 2020). Not only do these models better represent the adult human cell types, but discrepancies in our findings between the adult and hPSC-based models would be informative for understanding potential limitations of our ability to model adult behaviors in hPSC-derived cells.

Our preliminary analysis focused on finding perturbations that reveal cell-specific sensitivities in AP/CaT waveforms without generating arrhythmic phenotypes or loss of consistent pacing. However, these studies should be extended to identify pro-arrhythmic stimuli in both cell types. We observed that a number of early and delayed afterdepolarization events could be observed in our models by varying the pacing rates to model tachy/bradycardia conditions. This work could also be extended to include treatment with particular drugs in the setting of ion imbalances, which are known to increase the risk for arrythmias in patients (Haissaguerre et al., 2016). From this combinatorial analysis, we might identify conditions in which one cell type or the other is at higher risk for arrhythmia across a wide variety of conditions. Future work should also examine the extent to which hPSC-CMs and hPSC-PFs can model differences observed clinically in adult cell types, as these studies will help inform the use of these cells for further pharmacological testing.

## Author Contributions

DMG and NCD have conceptualized the study and written the manuscript with input from all authors. DMG, RD, JM, BS, JB, SS ad NCD have designed, performed and analyzed experiments. RD and ES have performed the mathematical modeling of the study. AM has generated the interactive data interface for the RNA-seq studies. SR has advice the study on the topic of Notch signaling and EP analysis.

## Acknowledgments

We thank all the members of the Dubois lab for constructive feedback, comments and ideas over the course of the project. We thank the Icahn School of Medicine at Mount Sinai (ISMMS) Stem Cell Engineering core (Samuele Marro, Erdene Balkinnyam), the ISMMS Microscopy facility (Deana Benson, Nikolaos Tzavaras, Glenn Doherty) and the ISMMS Flow Cytometry core (Jordi Ochando, Christopher Bare, Xuqiang Qiao) at the Icahn School of Medicine for their technical support. The work was funded by National Institutes of Health/National Heart Lung and Blood Institute (R01HL134956), the Additional Ventures Cures Collaborative, ISMMS seed funding and ISMMS Mindich Child Health and Development pilot funding to NCD. DMG was supported by an NRSF F31 Predoctoral Fellowship from the National Blood Lung and Heart Institute (5F31HL152612-02) and a T32 for training in Systems, Developmental Biology and Birth Defects from the Eunice Kennedy Shriver National Institute of Child Health and Development (HD075735). BS is supported by a pre-doctoral fellowship from the American Heart Association.

## Data Availability

RNA-seq data will be deposited in the NCBI Gene Expression Omnibus database.

---

**Supplementary Figure 1.**
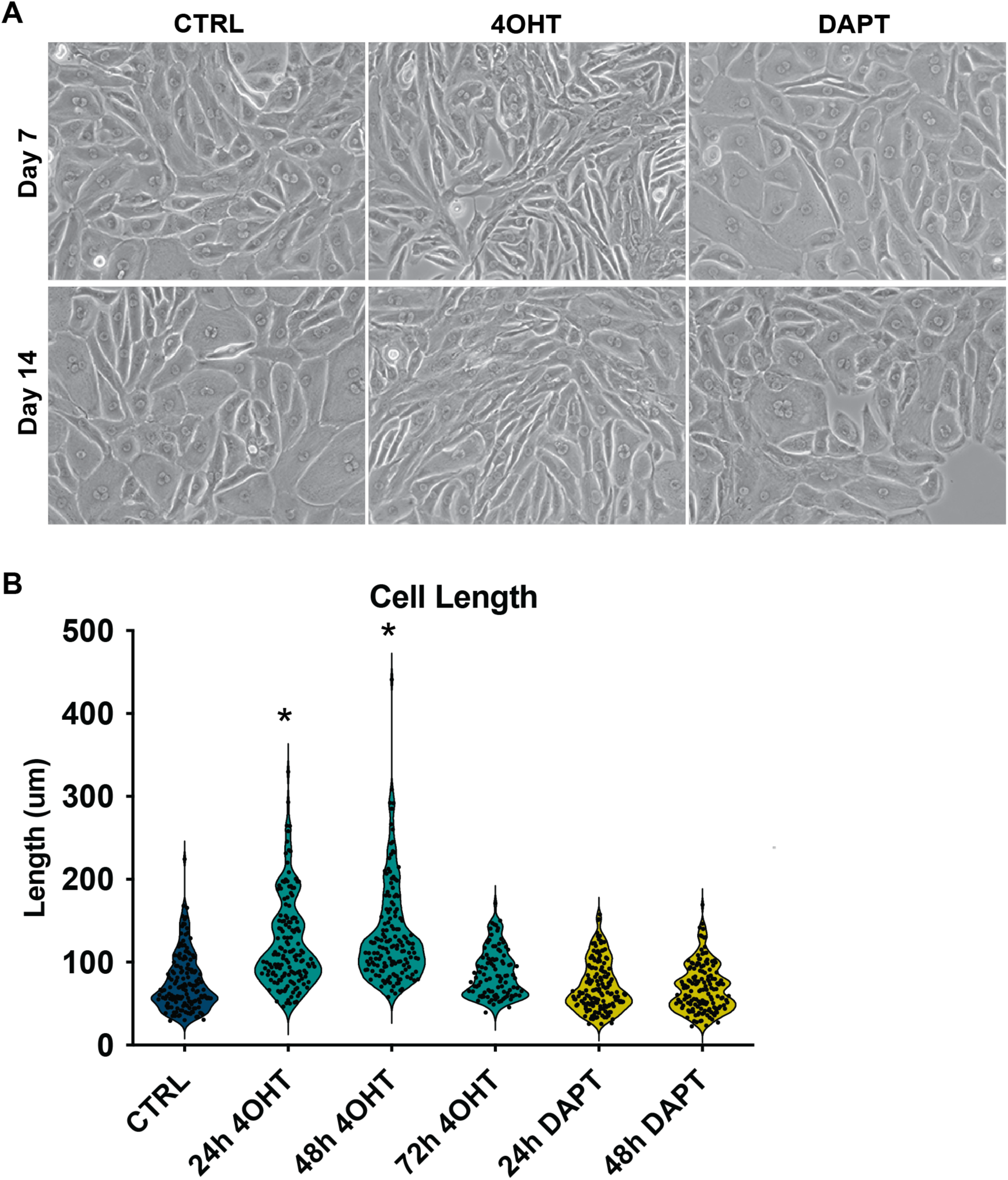
Acquisition of Elongated Morphology Following Notch Signaling Is Time-Dependent. **A)** Brightfield images showing gradual conversion toward elongated morphology over two weeks, following a 48h treatment with 4OHT. **B)** Calculation of cell length following treatment of 4OHT and DAPT for 24, 48, and 72 hours. Data collected from hPSC-CMs 28 days following 48h induction. * = p <0.05 according to students t-test

**Supplementary Figure 2.**
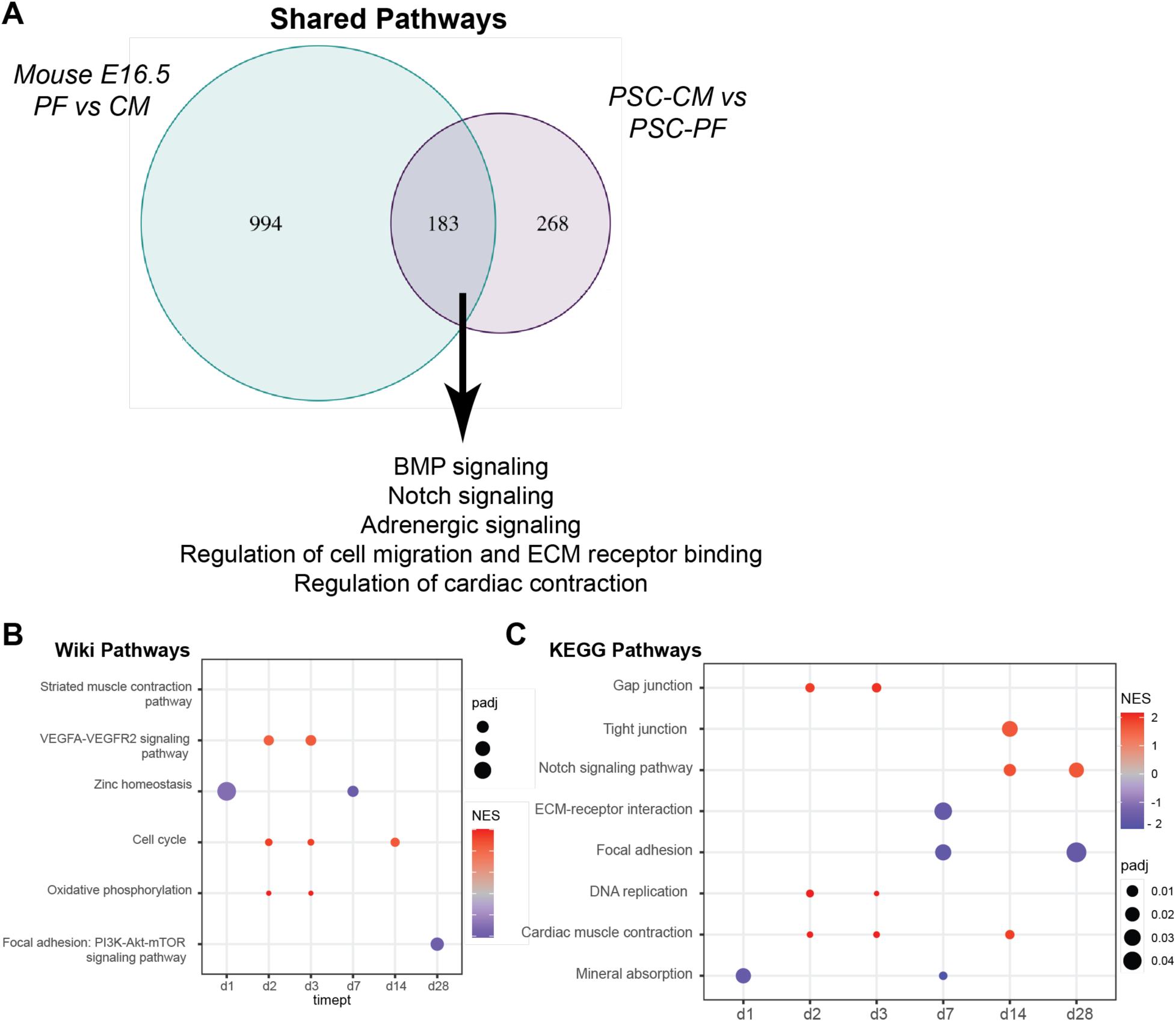
Transient Notch Activation in Pluripotent Stem Cell-Derived Cardiomyocytes Results in Transcriptomic Changes. **A)** Venn diagram demonstrating overlap of GO processes enriched in comparison between embryonic PFs and CMs from Goodyer et al and comparison between PSC-CMs and PSC-PFs. **B/C)** Bubble plot demonstrating enrichment of Wiki Pathways (B) and KEGG Pathways (C) at particular timepoints during conversion.

**Supplementary Figure 3.**
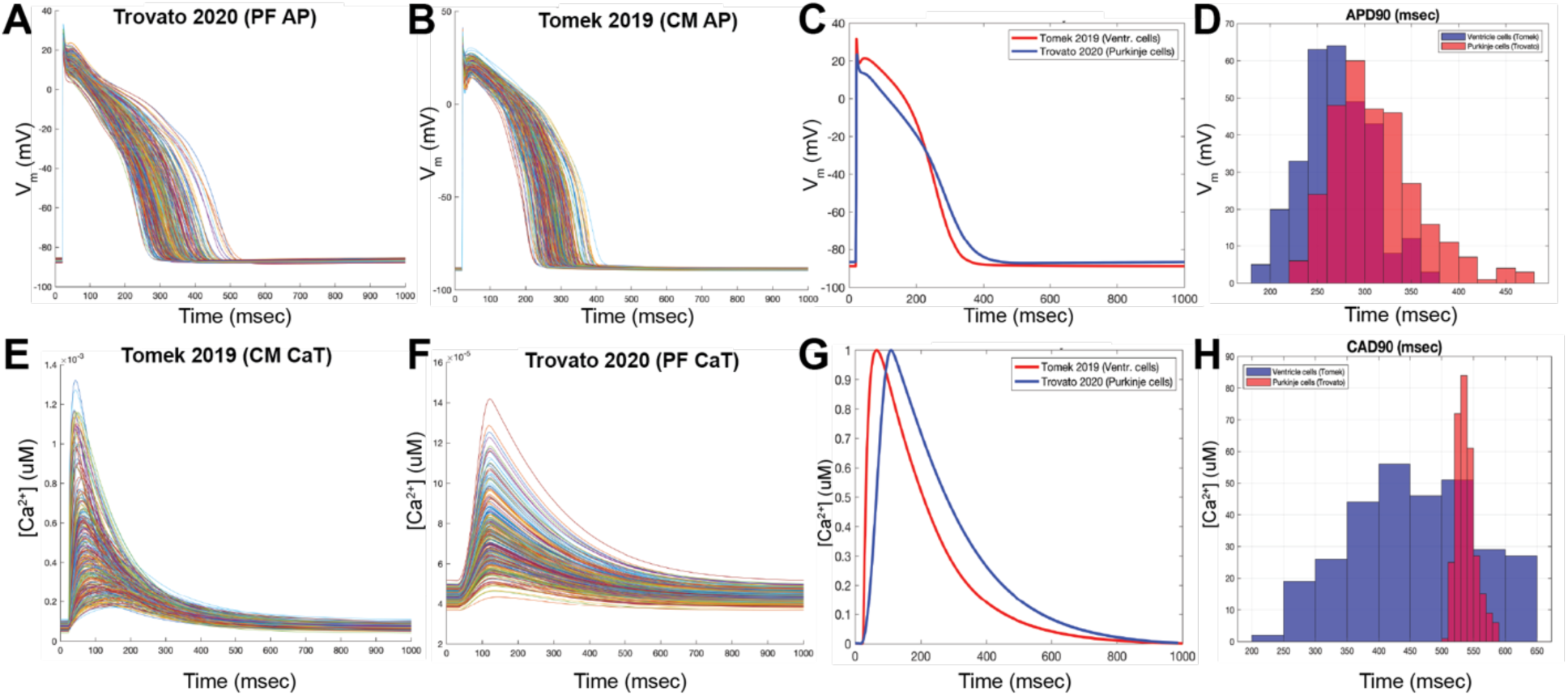
Direct Comparison of Human Post-Mortem Computational Models A/B) Multiple simulations of AP profiles using published models for **(A)** ventricular cardiomyocytes (VCM) and (**B)** Purkinje fibers (PFs). **C)** Mean AP profile for VCM (red) and PF (blue) models overlaid for comparison. **D)** Histogram demonstrating distribution of APD90 measurements after multiple rounds of simulation. **E/F)** Multiple simulations of calcium transient profiles using published models for **(E)** VCM and (**F)** PFs. **G)** Mean calcium transient for VCM (red) and PF (blue) models overlaid for comparison. **H)** Histogram demonstrating distribution of CAD90 measurements after multiple rounds of simulation.

**Supplementary File S1. In Silico Modeling of Drug Responses in Pluripotent Stem Cell Derived Cardiomyocytes and Purkinje Fibers (Extended)**

Files are organized in three folders. Raw = raw measurement of AP and CaT features. Delta = change in raw value of AP and CaT features compared to baseline untreated simulation. Delta Ratio = change in raw value of AP and CaT features expressed as a ratio of the change between PSC-PFs to PSC-CMs.

## MATERIALS AND METHODS

### Human pluripotent stem cell maintenance and differentiation

Human pluripotent stem cells (hPSCs) were maintained in E8 medium and passaged every 4 days onto matrigel-coated plates (Roche). The following hPSC lines were used in the study: *Rosa26;NICD-ER* line hESC line (generated and generously shared with us by Dr. Paul Gadue) and MSN02-4 line (human induced PSC generated at the Icahn School of Medicine)(Ditadi et al., 2015; Schaniel et al., 2021). On Day 0 (start of differentiation) hPSCs were treated with 1 mg/ml Collagenase B (Roche) for one hour, or until cells dissociated from plates, to generate embryoid bodies (EBs). Cells were collected and centrifuged at 300 rcf for 3 min, and resuspended as small clusters of 50–100 cells in differentiation medium containing RPMI (Gibco), 2 mmol/L L-glutamine (Invitrogen), 4×104 monothioglycerol (MTG, Sigma-Aldrich), 50 μg/ml ascorbic acid (Sigma-Aldrich). Differentiation medium was supplemented with 2 ng/ml BMP4 and 3 μmol Thiazovivin (Milipore). EBs were cultured in 6 cm dishes (USA Scientific) at 37°C in 5% CO2, 5% O2, and 90% N2. On Day 1, the medium was changed to differentiation medium supplemented with 30 ng/ml BMP4 (R&D Systems) and 30 ng/ml Activin A (R&D Systems), 5 ng/ml bFGF (R&D Systems) and 1 μmol Thiazovivin (Milipore). On Day 3, EBs were harvested and washed once with DMEM (Gibco). Medium was changed to differentiation medium supplemented with 5 ng/ml VEGF (R&D Systems) and 5 μmol/L XAV (Stemgent). On Day 5, medium was changed to differentiation medium supplemented with 5 ng/ml VEGF (R&D Systems). After Day 8, medium was changed every 3-4 days to differentiation medium without supplements for the remainder of differentiation.

### Cell dissociation, fluorescence-activated cell sorting (FACS), and plating for Notch perturbation

#### Cell Dissociation

EBs were dissociated between day 20-30 of differentiation. EBs were incubated overnight with 0.6 mg/ml Collagenase Type II (Worthington) at 37C. Dis-sociated cells were harvested and washed with Wash medium (DMEM, 0.1% BSA) + 1mg/ml DNase (VWR) twice and centrifuged at 300 g for 3 mins. Cells were resuspended in differentiation medium supplemented with 1 μM Thiazovivin (Millipore), filtered and counted using a hemacytometer. hPSC-CMs were plated onto matrigel-coated plates at appropriate densities per downstream application (12 well plate – 500,000 cells/well; 22X22 mm coverslips for immunofluorescence or AP/CaT analysis – 30,000 cells/cover slip; 48 well multi-electrode array – 60,000 cells/well). Media was changed the following day to remove Thiazovivin from media and maintained in differentiation medium. Medium was changed every 2 days.

#### FACS

Apart from scSeq experiments, all experiments were conducted on sorted (SIRPA+/CD90-) CMs. Dissociated cells were resuspended for 30 minutes on ice in differentiation medium containing antibodies

#### Notch signaling perturbation

Notch signaling was activated/inhibited 7 days after plating of cells as follows: 1 uM 4OHT (Sigma-Aldrich) and 10 uM DAPT (Sigma-Aldrich) were added to differentiation medium and added to cells for 48 hours. Medium was then changed to differentiation medium and maintained for up to

### Multi-electrode array analysis

Sorted SIRPA+/CD90-CMs were plated at 60,000 cells per well in a CytoView 48-well multielectrode array (MEA) plate (Axion Biosystems). One week after plating, Notch signaling was perturbed in these cells as described above. Cardiac Beat Detector measurement parameters were set at 150 μV detection threshold, 250 ms min. beat period, 5s max. beat period, polynomial regression FPD detection method, 70 ms post-spike detection holdoff, 50 ms pre-spike detection holdoff, 1s match post search duration with limit FPD search, and 10 running average beat count used for FPD detection. Axis software (Axion Biosystems) was utilized for downstream analysis following manufacturer’s instructions.

### Intracellular calcium transient (CaT) and action potential (AP) analysis

FACS-isolated SIRPA+CD90-hPSC-CMs were plated at 30,000 cells/coverslip (Sarstedt). 7 days post-plating, Notch signaling was perturbed as previously described for a period of 48 hours, then maintained in differentiation medium for 28 days. AP and CaT measurements were performed using voltage and Ca2+ sensitive dyes. Briefly, coverslips were incubated in 1X Tyrodes’ buffer (140 mM NaCl, 5.4 mM KCl, 10mM HEPES, 1 mM NaH2PO4, 1 mM MgCl2, 10 mM Glucose and 1.8 mM CaCl2, calibrated to pH=7.4) containing 10 mM Calbryte-630 AM (AAT Bioquest) and Fluovolt dyes (Invitrogen – dilution 1:1000 Fluovolt dye, 1:100 Powerload concentrate) in order to obtain paired AP and CaT recordings from a single coverslip. Coverslips were incubated in dye solution for 30 minutes at 37C. Coverslips were then transferred to custom cassette insert and incubated in 1X Tyrode’s buffer with 10 mM blebbistatin for 5 minutes, then loaded onto microscope for measurement. Tyrodes’ buffer was perfused throughout measurement to maintain cells at 37C. Cells were paced at 0.5 Hz using a MyoPacer Field Stimulator. 10-15 cells were recorded per coverslip across three biological replicates, resulting in a total of 30-45 cells per condition. For dofetilide treatments, a baseline recording of 10-15 cells was collected in standard Tyrodes’ buffer after which time the coverslip was perfused with a Tyrode’s solution containing 1,10, 25 and 50 nM dofetilide for 3-5 minutes and then re-recorded. Working solutions were diluted in Tyrode’s buffer from a 1 mM dofetilide solution reconstituted in DMSO.

Recordings of fluorescence flux in line scan were taken on an LSM5 exciter microscope using Zeiss ZEN 2009 software. Image files were analyzed using custom MATLAB scripts to extract features from individual waveforms for quantification.

### Immunofluorescence analysis and image processing

Cells plated on 22×22mm coverslips or tissue cryosections were rinsed with PBS and fixed with 4% Paraformaldehyde for 10 mins at room temperature. The edges of the coverslips were marked with a hydrophobic pen. Fixed cells were blocked with Saponin buffer (PBS, 0.5% Saponin, 0.1% BSA) for one hour at room temperature, incubated in primary antibody solution in Saponin buffer overnight at 4°C or for one hour at room temperature. Cells were rinsed three times with Saponin buffer, resuspended in secondary antibody diluted in Saponin buffer and incubated for one hour at room temperature. Cells were rinsed three times with Saponin buffer. Coverslips were mounted onto slides with nPG Antifade mounting medium (nPG stock : Glycerol : 10X PBS in the ratio 1:90:10) and edges were sealed with nail polish (VWR).

#### Primary antibody dilutions

Anti-α-Actinin (Sigma-Aldrich, 1:300), Anti-Cardiac Troponin T (Abcam, 1:400), Anti-Nav1.5 (Alomone Labs, 1:50), Anti-CX40 (Thermo Fisher, 1:250), Anti-CACNA2D2 (Alomone Labs, 1:100).

#### Secondary antibody dilutions

Alexa Fluor dyes (Jackson Immunoresearch) were diluted in Saponin buffer at 1:500.

#### Image acquisition and processing

All images were acquired on a Leica DM5500 inverted microscope and processed for brightness and contrast using ImageJ (Schindelin et al., 2012).

### cDNA generation and real-time qPCR

∼150,000 FACS isolated SIRPA+/CD90-CMs were collected, spun down at 300g for 3 minutes and snap frozen on dry ice. RNA was isolated from cell pellets using the Quick-RNA kit (Zymo Research). cDNA was generated using reverse transcription with the Quanta qScript kit (Quanta bio). Quantitative PCR was carried out on an Applied Biosystems Step One Plus using ABI SYBR Green reagents. Expression levels were normalized to TATA-binding protein (TBP) and genomic DNA was used for quantification of absolute expression levels, as previously described (Dubois et al., 2011).

### Bulk RNA sequencing

RNA was extracted from FACS-isolated SIRPA+CD90-hPSC-CMs using the Quick RNA Micro kit (Zymo Research). 500 ng of RNA per sample was used for Automated RiboZero Gold library prep and sequenced using a S2 100 cycle flow cell. FASTQ files were aligned to a reference transcriptome model generated from the human genome hg38 using STAR default parameters (Bray et al., 2016). The average number of reads successfully aligned across samples was ∼37 million. Count matrices were generated using featureCounts with default parameters. DESeq2 was used to normalize read counts and determine differentially expressed genes with p<0.05. ClusterProfiler was used to perform gene set enrichment analysis for GO/KEGG/WikiPathways that were upregulated or downregulated with p<0.05. DESeq2 and/or custom R scripts were used to generate heatmaps.

### dyn-EHT Fabrication, Mechanical Loading and Culture

Fabrication of dyn-EHTs was based on previously established methods where EHTs can be fabricated around polydimethylsolixane (PDMS) strips of known bending stiffness (Bliley et al. Science Translational Medicine 2020). Briefly, EHTs were cast around a PDMS strip (260 um thickness). Tissue components consisted of 18.75 million total cells per mL with a 90:10 ratio of cardiomyocytes:cardiac fibroblasts in a 1 mg/mL type I collagen (354249, Corning), 1.7 mg/mL Matrigel (354263, Corning) mixture. EHTs were cultured around the PDMS strips for 14 days to allow tissue compaction around the strip. At day 14, EHTs were removed from constrained PDMS wells and placed into dynamic culture for an additional 14 days. Media was changed to Cardiac Tissue Media consisting of RPMI 1640 supplemented with 1% knockout serum replacement (KOSR;10828028, Gibco) every 2-3 days during the experiment.

### Tissue contractility assay and measurements

To determine tissue contractile force, EHTs were viewed from the side in order to observe PDMS strip bending based on previously described methods (Bliley et al. Science Translational Medicine 2020). Briefly, the PDMS strip bending during EHT contraction was imaged with a Prime 95b camera mounted on a Nikon SMZ1500 stereomicroscope. During the contractility assay, tissues were maintained in Tyrode’s solution (T2145, Sigma-Aldrich) on a custom-heated stage at 37C +/- 1C. A custom-made MATLAB program (zenodo citation-from STM paper) was created to automatically calculate the change in length of the tissue from contraction videos. Look-up tables relating tissue length to force required to induce PDMS strip bending.

### Human pluripotent stem cell-derived Purkinje fiber computational model development – scaling of Paci et al model conductance through bulk RNA sequencing data

To simulate hPSC-PF behavior *in silico*, we scaled the original conductance for individual channels reported by Paci et al. 2018 for ventricular CMs and implemented the new parameters into the original Paci model (Paci et al., 2018). To scale conductance for individual ion channels, we used the results of our bulk RNA sequencing data comparing hPSC-CMs and hPSC-PFs. First, we assumed that the expression of ion channels of interest directly correlates with their ion channel densities (G_coi_, conductance of interest). Then, we extracted expression values for all the genes known to affect the following ion channels, pumps, exchangers (genes correlated): I_Na_ (SCN5A, SCN9A), I_K1_ (KCNJ2, KCNJ12), I_Kr_ (KCNH2), I_Ks_ (KCNQ1, KCNE1), I_to_ (KCND2, KCND3, KCNA4, KCNA7, KCNA7), I_CaL_(CACNA1S, CACNA1D, CACNA1B, CACNA1I, CACNA1G, CACNA1H, CACNA1A, CACNA1E, CACNA1F, CACNA1C, CACNA2D1,CACNB2), I_NaCa_ (SLC8A1), I_NaK_ (ATP1A1), I_RyR_ (RYR2), I_pCa_ (ATP2B4), I_up_ (ATP2A2), I_funny_ (HCN2, HCN4).

For ion currents with multiple genes, gene expression values were summed and the final expression values for each cell type were averaged across 3 biological samples from each group. hPSC-PF mean values for each ion channel were then divided by the expression of the equivalent ion channel expression of hPSC-CMs to create scaling factor for each ion current (PK/CM ratio). Finally, we modified the original conductances of the Paci model hPSC-CMs by multiplying each conductance value with its equivalent scaling factor.

### Cardiac cellular quantitative systems pharmacology modeling of drug effects

Both the original Paci hPSC-CM and hPSC-PF scaled models were used to simulate the effects of different drugs (e.g., dofetilide, nifedipine, amiodarone) on APs and CaT. Ion current inhibition by these drugs was simulated with a pore block model. With this approach, the conductance (G) of target ionic currents is scaled based on drug concentration ([C]), the IC_50_ value, and the Hill coefficient (H) that together describe how the drug blocks each current, for example:

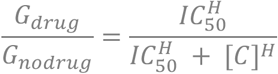

All simulation results were obtained during steady-state pacing at 1.5 Hz. An electrical stimulus current (1 ms duration, 40 μA/cm^2^ amplitude) was applied repeatedly to induce action potentials. A series of 200 consecutive stimuli were delivered to each cell/model, which usually caused cells to reach steady-state, meaning that consecutive action potentials were identical. When steady-state was reached, we quantified APD as the interval from the AP upstroke (maximal rate of rise) until membrane voltage decreased by 90% from peak level to resting level. Drug-induced AP prolongation was calculated as ΔAPD (i.e., drug-treated minus untreated cells).

### Statistical Analyses

Data are presented as mean ± standard deviation. Statistical significance between two groups was determined using two-sided unpaired or paired Student’s t-tests for parametric data, or Mann-Whiney test for non-parametric data as appropriate using GraphPad Prism 9. Statistical tests performed are included in the figure legends. For RNAseq analysis, statistical tests were performed using DESeq2/ClusterProfiler tools as outlined previously.

## REFERENCES

Árpádffy-Lovas, T., Husti, Z., Baczkó, I., Varró, A. and Virág, L. (2021). Different effects of amiodarone and dofetilide on the dispersion of repolarization between well-coupled ventricular and Purkinje fibers1. Can. J. Physiol. Pharmacol. 99, 48–55.

Baruteau, A.-E., Probst, V. and Abriel, H. (2015). Inherited progressive cardiac conduction disorders. Curr Opin Cardiol 30, 33–39.

Bray, N. L., Pimentel, H., Melsted, P. and Pachter, L. (2016). Near-optimal probabilistic RNA-seq quantification. Nat Biotechnol 34, 525–527.

Bruneau, B. G. and Srivastava, D. (2014). Congenital Heart Disease. Circulation Research 114, 598–599.

Burridge, P. W., Matsa, E., Shukla, P., Lin, Z. C., Churko, J. M., Ebert, A. D., Lan, F., Diecke, S., Huber, B., Mordwinkin, N. M., et al. (2014). Chemically defined generation of human cardiomyocytes. Nat Methods 11, 855–860.

Calderon, D., Bardot, E. and Dubois, N. (2016). Probing early heart development to instruct stem cell differentiation strategies. Dev Dyn 245, 1130–1144.

Chadwick, K., Wang, L., Li, L., Menendez, P., Murdoch, B., Rouleau, A. and Bhatia, M. (2003). Cytokines and BMP-4 promote hematopoietic differentiation of human embryonic stem cells. Blood 102, 906–915.

D’Amato, G., Luxán, G., del Monte-Nieto, G., Martínez-Poveda, B., Torroja, C., Walter, W., Bochter, M. S., Benedito, R., Cole, S., Martinez, F., et al. (2016a). Sequential Notch activation regulates ventricular chamber development. Nat Cell Biol 18, 7–20.

D’Amato, G., Luxán, G. and de la Pompa, J. L. (2016b). Notch signalling in ventricular chamber development and cardiomyopathy. FEBS J 283, 4223–4237.

Ditadi, A., Sturgeon, C. M., Tober, J., Awong, G., Kennedy, M., Yzaguirre, A. D., Azzola, L., Ng, E. S., Stanley, E. G., French, D. L., et al. (2015). Human definitive haemogenic endothelium and arterial vascular endothelium represent distinct lineages. Nat Cell Biol 17, 580–591.

Dubois, N. C., Craft, A. M., Sharma, P., Elliott, D. A., Stanley, E. G., Elefanty, A. G., Gramolini, A. and Keller, G. (2011). SIRPA is a specific cell-surface marker for isolating cardiomyocytes derived from human pluripotent stem cells. Nat Biotechnol 29, 1011– 1018.

Feulner, L., van Vliet, P. P., Puceat, M. and Andelfinger, G. (2022). Endocardial Regulation of Cardiac Development. J Cardiovasc Dev Dis 9, 122.

Goodyer, W. R., Beyersdorf, B. M., Paik, D. T., Tian, L., Li, G., Buikema, J. W., Chirikian, O., Choi, S., Venkatraman, S., Adams, E. L., et al. (2019). Transcriptomic Profiling of the Developing Cardiac Conduction System at Single-Cell Resolution. Circ Res 125, 379– 397.

Grego-Bessa, J., Luna-Zurita, L., del Monte, G., Bolós, V., Melgar, P., Arandilla, A., Garratt, A. N., Zang, H., Mukouyama, Y.-S., Chen, H., et al. (2007). Notch signaling is essential for ventricular chamber development. Dev Cell 12, 415–429.

Haissaguerre, M., Vigmond, E., Stuyvers, B., Hocini, M. and Bernus, O. (2016). Ventricular arrhythmias and the His-Purkinje system. Nat Rev Cardiol 13, 155–166.

Han, W., Wang, Z. and Nattel, S. (2000). A comparison of transient outward currents in canine cardiac Purkinje cells and ventricular myocytes. American Journal of Physiology-Heart and Circulatory Physiology 279, H466–H474.

Iyer, D., Gambardella, L., Bernard, W. G., Serrano, F., Mascetti, V. L., Pedersen, R. A., Talasila, A. and Sinha, S. (2015a). Robust derivation of epicardium and its differentiated smooth muscle cell progeny from human pluripotent stem cells. Development 142, 1528–1541.

Iyer, V., Roman-Campos, D., Sampson, K. J., Kang, G., Fishman, G. I. and Kass, R. S. (2015b). Purkinje Cells as Sources of Arrhythmias in Long QT Syndrome Type 3. Sci Rep 5, 13287.

Kattman, S. J., Witty, A. D., Gagliardi, M., Dubois, N. C., Niapour, M., Hotta, A., Ellis, J. and Keller, G. (2011). Stage-specific optimization of activin/nodal and BMP signaling promotes cardiac differentiation of mouse and human pluripotent stem cell lines. Cell Stem Cell 8, 228–240.

Kaufman, D. S., Hanson, E. T., Lewis, R. L., Auerbach, R. and Thomson, J. A. (2001). Hematopoietic colony-forming cells derived from human embryonic stem cells. Proc Natl Acad Sci U S A 98, 10716–10721.

Khandekar, A., Springer, S., Wang, W., Hicks, S., Weinheimer, C., Diaz-Trelles, R., Nerbonne, J. M. and Rentschler, S. (2016). Notch-Mediated Epigenetic Regulation of Voltage-Gated Potassium Currents. Circ Res 119, 1324–1338.

Kristóf, A., Husti, Z., Koncz, I., Kohajda, Z., Szél, T., Juhász, V., Biliczki, P., Jost, N., Baczkó, I., Papp, J., et al. (2012). Diclofenac Prolongs Repolarization in Ventricular Muscle with Impaired Repolarization Reserve. PloS one 7, e53255.

Lee, J. H., Protze, S. I., Laksman, Z., Backx, P. H. and Keller, G. M. (2017). Human Pluripotent Stem Cell-Derived Atrial and Ventricular Cardiomyocytes Develop from Distinct Mesoderm Populations. Cell Stem Cell 21, 179–194.e4.

Li, Y., Tian, X., Zhao, H., He, L., Zhang, S., Huang, X., Zhang, H., Miquerol, L. and Zhou, B. (2018). Genetic targeting of Purkinje fibres by Sema3a-CreERT2. Sci Rep 8, 2382.

Lian, X., Hsiao, C., Wilson, G., Zhu, K., Hazeltine, L. B., Azarin, S. M., Raval, K. K., Zhang, J., Kamp, T. J. and Palecek, S. P. (2012). Robust cardiomyocyte differentiation from human pluripotent stem cells via temporal modulation of canonical Wnt signaling. Proceedings of the National Academy of Sciences 109, E1848–E1857.

Lian, X., Zhang, J., Azarin, S. M., Zhu, K., Hazeltine, L. B., Bao, X., Hsiao, C., Kamp, T. J. and Palecek, S. P. (2013). Directed cardiomyocyte differentiation from human pluripotent stem cells by modulating Wnt/β-catenin signaling under fully defined conditions. Nature protocols 8, 162–75.

Lin, W. and Li, D. (2018). Zinc and Zinc Transporters: Novel Regulators of Ventricular Myocardial Development. Pediatr Cardiol 39, 1042–1051.

Lin, W., Li, D., Cheng, L., Li, L., Liu, F., Hand, N. J., Epstein, J. A. and Rader, D. J. (2018). Zinc transporter Slc39a8 is essential for cardiac ventricular compaction. J Clin Invest 128, 826–833.

Luxán, G., D’Amato, G. and de la Pompa, J. L. (2016). Intercellular Signaling in Cardiac Development and Disease: The NOTCH pathway. In Etiology and Morphogenesis of Congenital Heart Disease: From Gene Function and Cellular Interaction to Morphology (ed. Nakanishi, T.), Markwald, R. R.), Baldwin, H. S.), Keller, B. B.), Srivastava, D.), and Yamagishi, H.), p. Tokyo: Springer.

Maass, K., Shekhar, A., Lu, J., Kang, G., See, F., Kim, E. E., Delgado, C., Shen, S., Cohen, L. and Fishman, G. I. (2015). Isolation and characterization of embryonic stem cell-derived cardiac Purkinje cells. Stem Cells 33, 1102–1112.

MacGrogan, D., Nus, M. and de la Pompa, J. L. (2010). Notch signaling in cardiac development and disease. Curr Top Dev Biol 92, 333–365.

Mikryukov, A. A., Mazine, A., Wei, B., Yang, D., Miao, Y., Gu, M. and Keller, G. M. (2021). BMP10 Signaling Promotes the Development of Endocardial Cells from Human Pluripotent Stem Cell-Derived Cardiovascular Progenitors. Cell Stem Cell 28, 96–111.e7.

Miquerol, L., Moreno-Rascon, N., Beyer, S., Dupays, L., Meilhac, S. M., Buckingham, M. E., Franco, D. and Kelly, R. G. (2010). Biphasic development of the mammalian ventricular conduction system. Circ Res 107, 153–161.

Mohan, R. A., Boukens, B. J. and Christoffels, V. M. (2018). Developmental Origin of the Cardiac Conduction System: Insight from Lineage Tracing. Pediatr Cardiol 39, 1107– 1114.

Murry, C. E. and Keller, G. (2008). Differentiation of embryonic stem cells to clinically relevant populations: lessons from embryonic development. Cell 132, 661–680.

Nakanishi, N., Takahashi, T., Ogata, T., Adachi, A., Imoto-Tsubakimoto, H., Ueyama, T. and Matsubara, H. (2012). PARM-1 promotes cardiomyogenic differentiation through regulating the BMP/Smad signaling pathway. Biochem Biophys Res Commun 428, 500– 505.

Paci, M., Pölönen, R.-P., Cori, D., Penttinen, K., Aalto-Setälä, K., Severi, S. and Hyttinen, J. (2018). Automatic Optimization of an in Silico Model of Human iPSC Derived Cardiomyocytes Recapitulating Calcium Handling Abnormalities. Front Physiol 9, 709.

Park, D. S. and Fishman, G. I. (2017). Development and Function of the Cardiac Conduction System in Health and Disease. J Cardiovasc Dev Dis 4, 7.

Prodan, N., Ershad, F., Reyes-Alcaraz, A., Li, L., Mistretta, B., Gonzalez, L., Rao, Z., Yu, C., Gunaratne, P. H., Li, N., et al. (2022). Direct reprogramming of cardiomyocytes into cardiac Purkinje-like cells. iScience 25, 105402.

Protze, S. I., Liu, J., Nussinovitch, U., Ohana, L., Backx, P. H., Gepstein, L. and Keller, G. M. (2017). Sinoatrial node cardiomyocytes derived from human pluripotent cells function as a biological pacemaker. Nat Biotechnol 35, 56–68.

Rentschler, S., Yen, A. H., Lu, J., Petrenko, N. B., Lu, M. M., Manderfield, L. J., Patel, V. V., Fishman, G. I. and Epstein, J. A. (2012). Myocardial Notch signaling reprograms cardiomyocytes to a conduction-like phenotype. Circulation 126, 1058–1066.

Schaniel, C., Dhanan, P., Hu, B., Xiong, Y., Raghunandan, T., Gonzalez, D. M., Dariolli, R., D’Souza, S. L., Yadaw, A. S., Hansen, J., et al. (2021). A library of induced pluripotent stem cells from clinically well-characterized, diverse healthy human individuals. Stem Cell Reports 16, 3036–3049.

Schindelin, J., Arganda-Carreras, I., Frise, E., Kaynig, V., Longair, M., Pietzsch, T., Preibisch, S., Rueden, C., Saalfeld, S., Schmid, B., et al. (2012). Fiji: an open-source platform for biological-image analysis. Nat Methods 9, 676–682.

Stankunas, K., Hang, C. T., Tsun, Z.-Y., Chen, H., Lee, N. V., Wu, J. I., Shang, C., Bayle, J. H., Shou, W., Iruela-Arispe, M. L., et al. (2008). Endocardial Brg1 represses ADAMTS1 to maintain the microenvironment for myocardial morphogenesis. Dev Cell 14, 298–311.

Terrar, D. A., Wilson, C. M., Graham, S. G., Bryant, S. M. and Heath, B. M. (2007). Comparison of guinea-pig ventricular myocytes and dog Purkinje fibres for in vitro assessment of drug-induced delayed repolarization. J Pharmacol Toxicol Methods 56, 171–185.

Tomek, J., Bueno-Orovio, A., Passini, E., Zhou, X., Minchole, A., Britton, O., Bartolucci, C., Severi, S., Shrier, A., Virag, L., et al. (2019). Development, calibration, and validation of a novel human ventricular myocyte model in health, disease, and drug block. Elife 8, e48890.

Trovato, C., Passini, E., Nagy, N., Varró, A., Abi-Gerges, N., Severi, S. and Rodriguez, B. (2020). Human Purkinje in silico model enables mechanistic investigations into automaticity and pro-arrhythmic abnormalities. J Mol Cell Cardiol 142, 24–38.

Tsai, S.-Y., Maass, K., Lu, J., Fishman, G. I., Chen, S. and Evans, T. (2015). Efficient Generation of Cardiac Purkinje Cells from ESCs by Activating cAMP Signaling. Stem Cell Reports 4, 1089–1102.

van Eif, V. W. W., Devalla, H. D., Boink, G. J. J. and Christoffels, V. M. (2018). Transcriptional regulation of the cardiac conduction system. Nat Rev Cardiol 15, 617–630.

van Weerd, J. H. and Christoffels, V. M. (2016). Regulation of Vertebrate Conduction System Development. In Etiology and Morphogenesis of Congenital Heart Disease: From Gene Function and Cellular Interaction to Morphology (ed. Nakanishi, T.), Markwald, R. R.), Baldwin, H. S.), Keller, B. B.), Srivastava, D.), and Yamagishi, H.), p. Tokyo: Springer.

Yang, L., Geng, Z., Nickel, T., Johnson, C., Gao, L., Dutton, J., Hou, C. and Zhang, J. (2016). Differentiation of Human Induced-Pluripotent Stem Cells into Smooth-Muscle Cells: Two Novel Protocols. PLoS One 11, e0147155.

Zhao, L., Ben-Yair, R., Burns, C. E. and Burns, C. G. (2019). Endocardial Notch Signaling Promotes Cardiomyocyte Proliferation in the Regenerating Zebrafish Heart through Wnt Pathway Antagonism. Cell Rep 26, 546–554.e5.

